# Long-read transcriptomic identification of synaptic adaptation to amyloid pathology in the *App^NL-G-F^* knock-in mouse model of the earliest phase of Alzheimer’s disease

**DOI:** 10.1101/2024.08.10.607237

**Authors:** Umran Yaman, Gareth Banks, Emil K. Gustavsson, Paige Mumford, Naciye Magusali, Orjona Stella Taso, Hannah Macpherson, Susana Carmona, Malgorzata Murray, Rasneer Sonia Bains, Hamish Forrest, Michelle Stewart, Connor Scott, Tatiana V. Lipina, Zhao Cheng, Anna L. Tierney, Richard D. Unwin, Juan A. Botia, Carlo Sala Frigerio, Sara E. Wells, John Hardy, Lilach Soreq, Frances K. Wiseman, Dervis A. Salih

## Abstract

Genome-wide association studies (GWAS) have identified a transcriptional network of Alzheimer’s disease (AD) risk genes that are primarily expressed in microglia and are associated with AD pathology. However, traditional short-read sequencers have limited our ability to fully characterize how GWAS variants exert their effects on gene expression regulation or alternative splicing in response to the pathology, particularly resulting in inaccurate detection of splicing. To address this gap, we utilized long-read RNA-sequencing (RNA-seq) in the *App^NL-G-F^* knock-in mouse model to identify changes in splicing and novel transcript isoforms in response to amyloid-β. We show that long-read RNA-seq can recapitulate the expected induction of microglial expressed risk genes such as *Trem2* in response to amyloid-β at 9 months of age associated with ageing-dependent deficiencies in spatial short-term memory in the *App^NL-G-F^* knock-in mice. Our results not only identified novel splicing events and transcript isoform abundance in genes associated with AD, but also revealed the complex regulation of gene expression through splicing in response to amyloid plaques. Surprisingly, the regulation of alternative splicing in response to amyloid was seen in genes previously not identified as AD risk genes, expressed in microglia, neurons and oligodendrocytes, and included genes such as *Syngr1* that modulate synaptic physiology. We saw alternative splicing in genes such as *Ctsa*, *Clta*, *Dennd2a*, *Irf9* and *Smad4* in mice in response to amyloid, and the orthologues of these genes also showed transcript usage changes in human AD brains. Our data suggests a model whereby induction of AD risk gene expression associated with microglial proliferation and activation is concomitant with alternative splicing in a different class of genes expressed by microglia and neurons, which act to adapt or preserve synaptic activity in response to amyloid-β during early stages of the disease. Our study provides new insights into the mechanisms and effects of the regulation of genes associated with amyloid pathology, which may ultimately enable better disease diagnosis, and improved tracking of disease progression. Additionally, our findings identify new therapeutic avenues for treatment of AD.

## Introduction

Late-onset Alzheimer’s disease (AD) is typically characterised by an extended asymptomatic phase marked by the progressive accumulation of amyloid and subsequently tau pathology. This pathological progression occurs across distinct cellular states, encompassing neurons, glia, oligodendrocytes, and the vascular system, culminating in neuronal death approximately two decades following disease onset^1^. Although the source of amyloid and tau are usually considered to be neuronal, a decade of genome-wide association studies (GWAS) revealed that most gene variants associated with AD risk were expressed by the innate immune system^2–5^. This indicates that microglia/monocytes likely play a substantial role in modifying the risk of AD through variants of genes such as Apolipoprotein E (*APOE*), Triggering Receptor Expressed on Myeloid Cells 2 (*TREM2),* Spi-1 Proto-Oncogene *(SPI1),* Phospholipase C Gamma 2 (*PLCG2*) and 2’-5’-Oligoadenylate Synthetase 1 (*OAS1)*^5,6^. Transcriptome studies in mouse models using microarrays, bulk short-read RNA-seq and single-cell RNA-seq have shown that amyloid plaques alone are sufficient to modify expression of these microglial risk genes and induce microglial changes including proliferation and activation, indicating these AD risk genes and microglial adaptations may act at early stages of disease in humans to impact the onset and progression of clinical symptoms^5,7,8^. Overt cognitive deficits in humans typically lead to diagnosis when a substantial level of neuronal loss occurs^9,10^. This loss starts approximately twenty years before the onset of symptoms.

The extent to which mouse models can faithfully recapitulate the intricate features of human AD has been a subject of ongoing debate. The majority of mouse models do not exhibit the amyloid-dependent tau pathology that leads to neuronal degeneration^11,12^ which has raised questions about their relevance for AD research^13–15^. As a result, it is widely recognised that AD mouse models, and particularly mouse models with amyloid pathology, encompass a range of phenotypes that predominantly reflect various aspects of the early prodromal stages of the disease^16,17^. These include crucial processes such as Amyloid Precursor Protein (APP) processing, amyloid-β plaque deposition, and microglial activation which are mediated by the induction of risk genes identified in human GWAS^18^. In order to advance our understanding, it becomes important to undertake a characterisation of the earliest molecular and cellular alterations in response to amyloid pathology, and how this relates to clinically relevant cognitive changes. Characterisation of early changes, such as isoform-level alterations, have significant implications not only for the early diagnosis of AD, but also for stratifying patients, a crucial step in emerging clinical trials. This elucidation of early changes is poised to enhance our comprehension of how subtle alterations manifest in disease progression, and how transcript isoforms might impact translation, ultimately facilitating the identification of potential therapeutic targets.

GWAS have identified approximately one hundred loci associated with AD^2–4^, but with many of the variants and single nucleotide polymorphisms (SNPs) identified, the protein coding sequence remains unaffected, leaving their mechanistic contributions to AD largely unknown. These variants may exert influence on gene regulation by modifying gene expression and splicing patterns^19–22^. Alternative transcript isoforms and splice-forms are pivotal mechanisms in gene expression, and errors in splicing can lead to regulatory dysfunction^23–28^. Alternative splicing plays a crucial role in protein function, particularly in neurological diseases^29,30^, and selection of specific mRNA isoforms is instrumental in driving various cellular pathways and functions^21,31,32^.

While previous studies employing short-read RNA-seq have identified microglial networks associated with amyloid pathology that are enriched with isoforms of GWAS candidate genes^33,34,33,34^, the current genome assembly does not fully encompass the diversity of genes and isoforms, nor accurate splicing patterns^35^. Long-read RNA-sequencing has emerged as a powerful tool for elucidating splicing and transcript isoform expression, as well as identifying novel exons and splicing patterns that were previously challenging to discern on a genome-wide scale using conventional methods. In our study, we employed long-read RNA-seq in the *App^NL-G-F^* knock-in mouse model alongside wildtype littermate controls to discern changes in transcript usage, splicing and novel transcript isoforms in response to amyloid at an age in which the model is starting to develop AD relevant cognitive changes. By adopting an isoform-centric approach, we offer unbiased insights into changes in altenative splicing events^36^. Previous gene-level expression analyses lack the sensitivity needed to detect possible changes at the transcript level, such as alterations in splicing. To overcome this limitation, we employed differential transcript usage analysis to identify additional shifts in gene expression in *App^NL-G-F^* knock-in mouse models compared to controls. Our results reaffirm the significance and intricacy of known AD risk genes while also revealing novel splicing and potential regulatory patterns in some of these AD risk genes and also in a series of other genes not previously linked to AD risk.

Our analysis also uncovers novel splicing events and transcript isoforms that were not previously discernible through gene-level and isoform-level expression analysis in genes previously not thought to contribute to AD risk, particularly of the genes integral to synaptic physiology. Our analysis identified differential isoform usage in genes linked to pre- and post-synaptic functions which might implicate an association with synaptic resilience in AD^37^. This approach enables more precise annotations of isoforms, contributing to a deeper understanding of protein diversity and specific patterns of isoform changes that contribute to the disease. Such insights may prove valuable in detecting the earliest stages of disease.

## Results

### Behavioural and sensory phenotyping of *App^NL-G-F/NL-G-F^* knock-in animals

To investigate transcript isoform usage and splicing in response to amyloid pathology we selected the commonly used *App^NL-G-F^* mice^38,39^ (Figure 1). We initially confirmed that amyloid accumulation was similar to previous studies with our housing conditions^38,39^ (Figures S1 and S2). To relate transcriptional changes to disease stage in this model of the earliest phases of AD, we determined if the accumulation of amyloid-β in the brain or changes to APP biology altered cognitive function, locomotor activity, sensory systems or body composition, in *App^NL-G-F/NL-G-F^* knock-in mice and colony-matched C57BL/6J between 2 and 9-months of age. All means and significance analyses are detailed in Tables S1, S2, S3 and S4.

**Figure 1.**
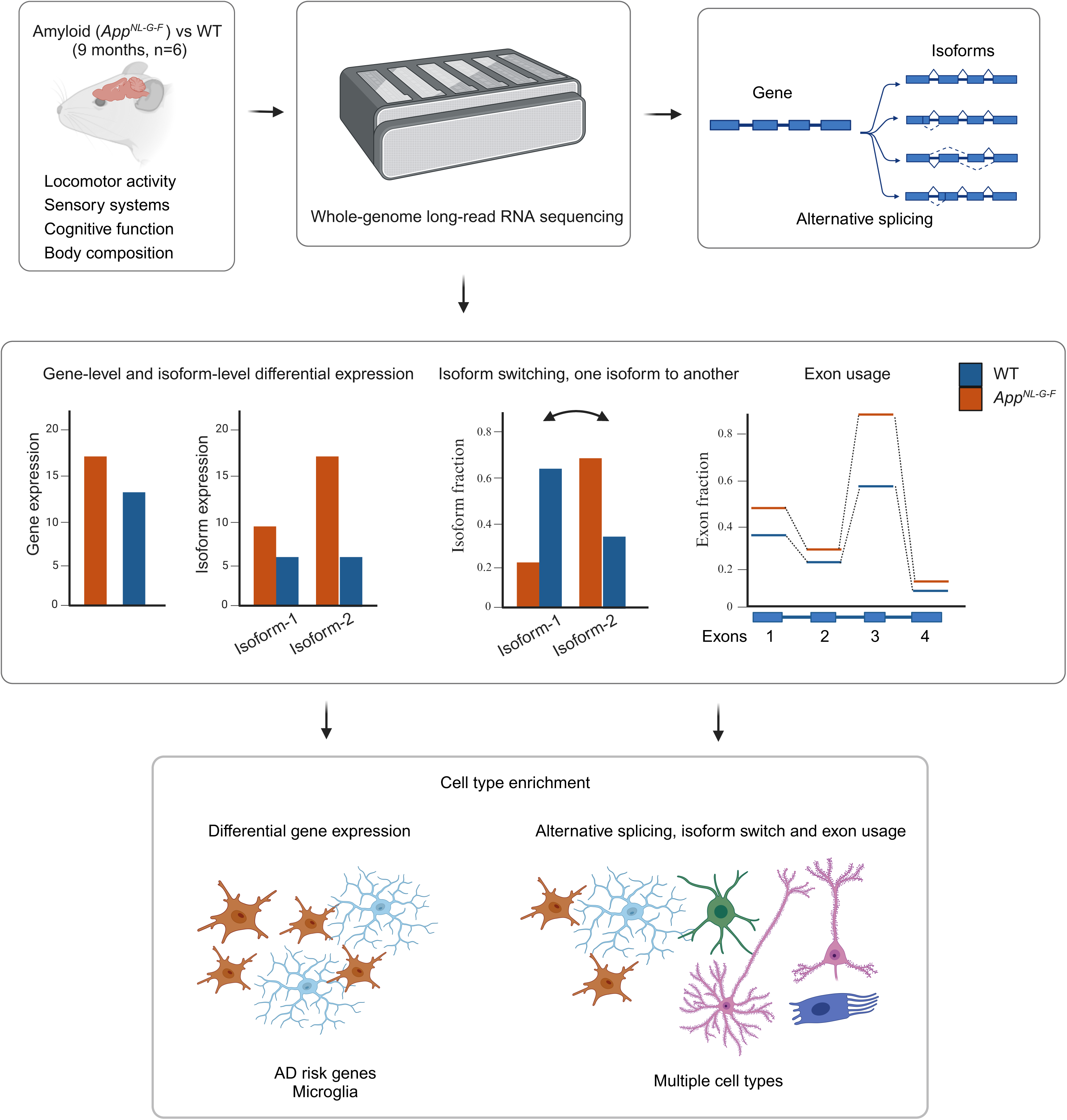
Experimental workflow to investigate transcript diversity genome-wide in response to amyloid pathology in the *App^NL-G-F^* mouse model of early AD using long-read RNA-seq. Phenotype severity was assessed for *App^NL-G-F^*knock-in versus control mice at 9 months of age by testing locomotor activity, sensory systems, cognitive function and body composition. Long-read RNA-seq was used to investigate gene-level expression, usage of individual transcript isoforms, isoform switching, exon usage, alternative splicing and alternative polyadenylation sequences.

Short-term learning and memory were tested using the novel object recognition task to assess recognition memory and the forced Y-maze to assess spatial memory. Testing using the novel object recognition task (at 14 and 34 weeks of age) showed no significant effects of genotype or any genotype interactions upon the parameters measured (Table S1). In contrast, testing using the forced Y-maze (at 12 and 33 weeks of age) demonstrated a significant interaction between genotype and age upon both the duration of time spent in the novel arm and the number of entries into the novel arm (genotype X age interaction; duration in novel arm: F[1,56]=6.8, p=0.012; number of entries into novel arm: F[1,56]=4.09, p=0.048). Pairwise analysis demonstrated that for both parameters at 33 weeks of age, *App^NL-G-F/NL-G-F^*animals showed a significantly reduced novelty preference ratio for both parameters compared with wildtype controls (duration in novel arm: p=0.007; number of entries in the novel arm: p=0.026). There was no significant difference between genotypes at 10 week of age (duration in novel arm: p=0.517; number of entries in the novel arm: p=0.629). We found no significant effects of genotype or genotype interactions in activity levels in either novel object recognition or the Y-maze, suggesting that activity changes were not confounds in these tests (Table S1). This indicates that a decline in spatial short-term memory occurs in the *App^NL-G-F^*knock-in mouse model between 12 and 33 weeks of age.

Locomotor activity was assessed by home cage monitoring in group housed conditions at 10 and 32 weeks of age^40^. Here, we found no effect of genotype upon activity levels in either the light or the dark phases of the light-dark cycle or at the transitions between the light and dark phases (Table S2).

*App* is expressed in a number of cell types throughout the body in addition to in neurons, thus changes to the biology of the gene in the *App^NL-G-F/NL-G-F^* may affect the structure and function of these cells. To determine if sensory inputs may contribute to the spatial memory deficit we observed at 33 weeks of age, we assessed the sensory systems using the Auditory Brainstem Response (ABR) to measure auditory function and Optical Coherence Tomography (OCT) to study the structure of the retina. ABR recordings (at 15 to 16 weeks of age) demonstrated no effect of genotype upon ABR thresholds (Table S3). Notably, the levels of evoked thresholds suggested that all animals could hear^41^.

*App* is expressed in retinal ganglion cells, and previous work has indicated that the biology of the eye may be perturbed in the *App^NL-G-F/NL-G-F^* model^42^, we undertook OCT scans of the retina (at 15-16 and 37-38 weeks of age). Visual inspection revealed no evidence of gross abnormalities in the retinal morphology of *App^NL-G-F/NL-G-F^* animals. However, analysis of the retina thickness demonstrated a significant impact of genotype, with *App^NL-G-F/NL-G-F^*animals having a thinner retina (F[1,55]=4.35; p=0.0416). Therefore we conducted segmentation analysis to analyse the size of the individual retinal layers. This analysis revealed significant impacts of genotype in the Inner Nuclear Layer (INL), the Ganglion Cell Layer (GCL) and the Outer Part of the Retina/Subretinal virtual space (OPR/Subretinal) area of the retina (see Table S3). Analysis of INL size demonstrated that *App^NL-G-F/NL-G-F^* animals have a significantly thinner INL than wildtypes (F[1,55]=4.59; p=0.036). Both the GCL and the OPR/Subretinal space showed significant interactions between age, sex and genotype (GCL: F[1,10]=6.53; p=0.028. OPR/Subretinal space: F[1,10]=13.32; p=0.005). Pairwise analysis of these retinal layers found no significant differences in male animals of either age or genotype. However, 37-38-week-old female *App^NL-G-F/NL-G-F^* animals had a significantly thinner CGL than wildtype controls (p=0.005). Consistent with this, female *App^NL-G-F^* animals showed a significant age-related increase in the OPR/Subretinal space size (15 weeks v 34 weeks: p=0.0084).

At 8, 17 and 39 weeks of age weight and lean mass were assessed by ECHO-MRI. We found no significant genotype effects on either weight or lean mass (Table S4).

### Gene-level expression changes confirm microglial proliferation and activation

To investigate the impact of amyloid pathology on transcription isoforms and splicing of gene expression, we performed long-read RNA-seq on total cortical RNA of *App^NL-G-F^*knock-in mice and littermate C57BL/6J controls at 9 months of age. Sequence reads were then aligned to the latest release of the Gencode mouse reference genome (GRCm39) with minimap2^43^. We analysed the long-read RNA-seq to determine differential gene-level expression, usage of individual transcript isoforms, exon switching, alternative splicing, alternative polyadenylation sequences and identifying changes to the protein coding sequence, using DESeq2^44^, DEXSeq^45^, PSI-Sigma^46^, IsoformSwitchAnalyzer^47^, and tappAS which is a part of Functional Iso-Transcriptomics (FIT) framework ^48^.

Our differential gene-level expression analysis using DESeq2 confirmed the expected gene-level changes at the age of 9 months in the cortex, as seen in previous studies with short read RNA-seq using the same mouse model^17^, and other mouse models with amyloid plaques^5,7,16^. Our analysis identified 178 differentially expressed genes (176 upregulated and 2 downregulated, |log_2_FC| > 0.5 and FDR < 0.05) in *App^NL-G-F^* knock-in versus control mice (Figure 2A; Table S5). These findings show a significant overlap with the amyloid-responsive microglia (ARM), also known as the disease-associated microglial (DAM) gene cluster identified in previous literature using single-cell RNA-seq analysis (Fisher’s exact test, p-value < 2.2e^-16^)^16^. The identified overlap includes many well-known orthologues of AD risk genes, such as *Trem2*, *Tyrobp* and *Ctsd*. Biological annotation of the differentially expressed genes highlighted the expected immune system associated processes in *App^NL-G-F^* mice (Figure 2B), reflecting the proliferation and activation of microglia. Network analysis to identify the co-expressed genes in response to amyloid produced a genetic network representing amyloid-responsive microglia, with Beta-2-Microglobulin (*B2m*), Complement Component 1, Q Subcomponent, B Chain (*C1qb*), Cathepsin D (*Ctsd*), Cathepsin Z (*Ctsz*), Granulin (*Grn*) and *Spi1* as hub genes, which are proposed to drive the observed transcriptional response to amyloid (Figure 2C; Table S6). This amyloid-responsive microglial network is similar to those we described previously by short-read RNA-seq and microarrays in other mouse models with amyloid plaques^5,7^.

**Figure 2.**
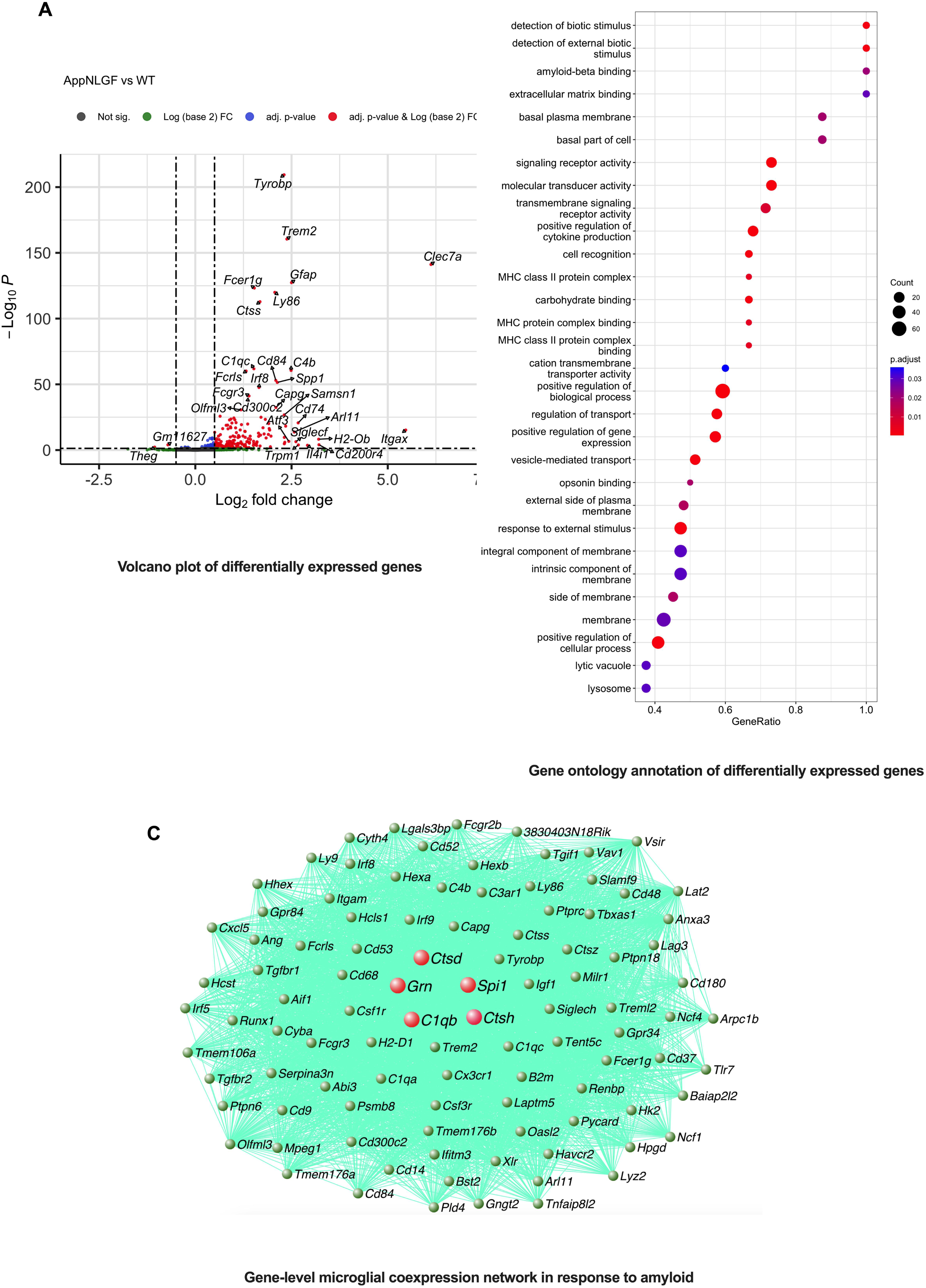
Gene-level differential expression analysis in *App^NL-G-F^* versus control mice at 9 months of age using long-read RNA-seq. (A) The differential expression analysis confirms the microglial response to amyloid plaques, with many genes associated with ARM/DAM (*e.g. Trem2, Tyrobp, Ctsd, C1qb*), assessed by DESeq2 (FDR < 0.05). The analysis was performed on a cohort of n = 6 mice for each genotype, with an equal representation of n = 3 males and 3 females per genotype. (B) Differentially expressed genes are enriched for immune cell activation pathways, indicating a microglial response to amyloid plaques. (C) The coexpression network analysis of gene-level expression using long-read RNA-seq reveals an overlap with the ARM/DAM generated using short-read sequencing data^16,17^.

We also investigated if changes in gene-expression were sex-dependent, but only a limited number of differentially expressed genes were observed to be dependent on sex at 9 months of age (6 differentially expressed genes; Table S7). These limited genes had no obvious enrichment for known biological annotations, and this indicates that sex-dependent changes in gene expression are limited at this relatively young age of 9 months.

### Isoform-level differential expression changes refine myeloid cell function

A number of isoforms, such as *Ccl6-201*, chemokine (C-C motif) ligand 6 (Figure 3A; Table S8), displayed substantial fold-expression changes at the transcript level as compared to whole gene expression level. Specific *Trem2* isoforms (ENSMUST00000113237.3, log_2_FC = 2.75, FDR = 3.41e^-118^, and ENSMUST00000024791.14, log_2_FC = 2.28, FDR = 6.41e^-42^), an *Apoe* isoform (ENSMUST00000174064.8), and a Glial Fibrillary Acidic Protein (*Gfap*) isoform (ENSMUST00000067444.9, log_2_FC = 2.60, FDR = 4.38e^-46^), amongst others, were differentialy expressed in the cortex of *App^NL-G-F^* mice compared to wildtype controls. This suggests that certain genes may exert effects via the transcript level, potentially driving specific microglial network functions, in response to the amyloid pathology (Figure S3). The hub isoforms represented by gene names in Figure S3 are likely to be major cellular drivers at the isoform-scale, which are not evident on the gene-level, suggesting specific isoforms of genes may contribute to the early pathological changes. Our isoform-level annotation via Gene Ontology (GO) terms revealed changes in chemokine activity, reduced endopeptidase activity, and signalling receptor regulation (Figure 3B). Our findings highlight the significance of isoform-level changes, which offer valuable insights into subtle yet significant cellular alterations that may go unnoticed at the gene-level. This observation emphasizes the importance of considering isoform-level dynamics to fully comprehend the molecular mechanisms underlying the response to pathology. Moreover, the enrichment of genes with altered expression within the broader innate immune response pathway suggests that these genes may possess specific roles or functions contributing to the response to pathology. These findings underscore the complex interplay between isoform-level regulation and the broader gene-level enrichment within the innate immune response pathway.

**Figure 3.**
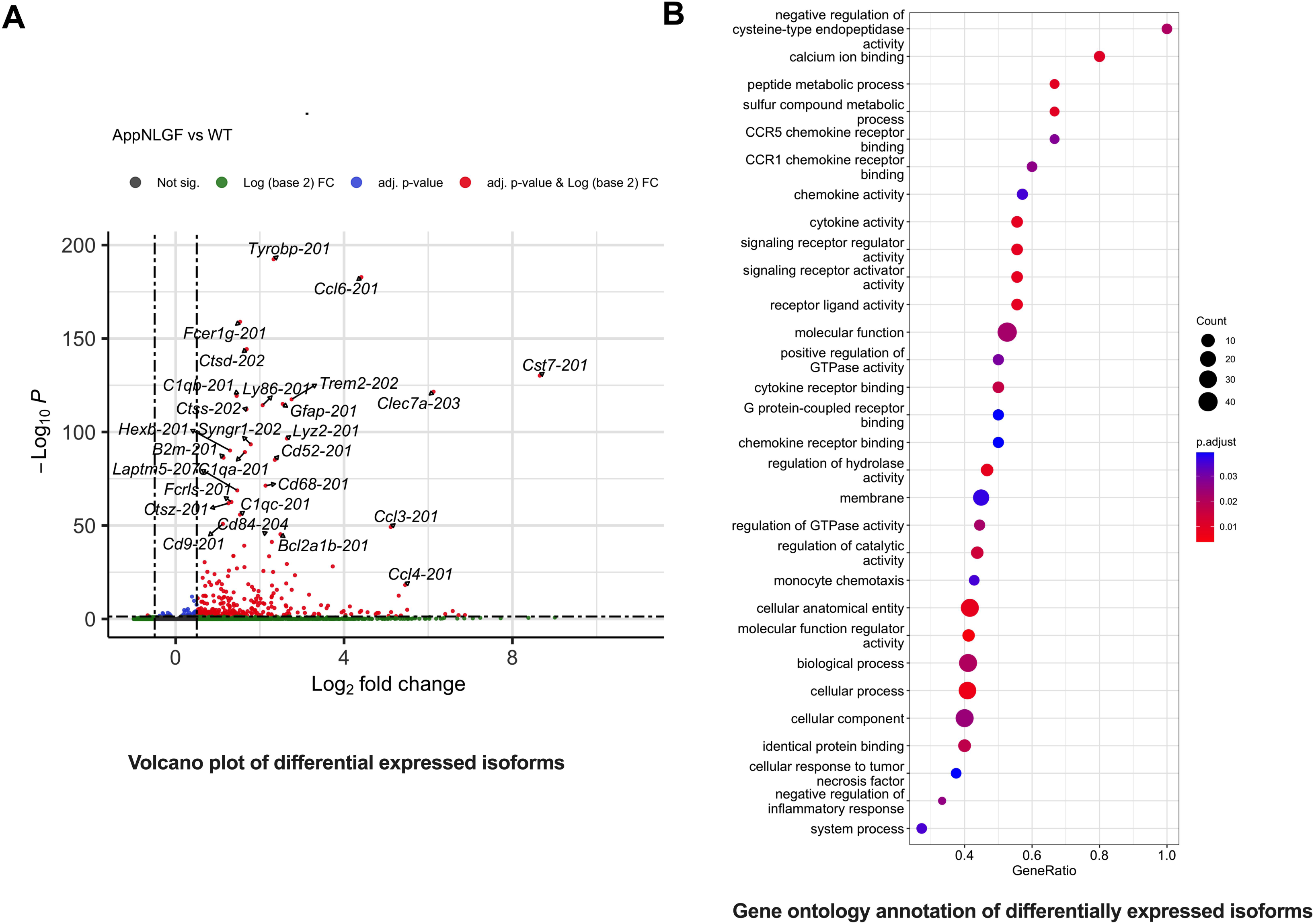
Transcript isoform-level expression analysis and enrichment of immune-related pathways. (A) Isoform-level expression analysis revealed that several transcripts, such as *Tyrobp-201*, *Ccl6-201*, exhibited greater fold-expression level changes at the transcript-level compared to the gene-level, assessed by DESeq2 (FDR < 0.05). This suggests strong preference bias of specific transcript isoforms in response to amyloid pathology (n = 6 mice per group). (B) Enrichment analysis showed prominent enrichment of cytokine activity, membrane transport and metabolism. GO enrichments highlight the importance of isoform-level changes in understanding the cellular response to pathology and underscore the interplay between isoform-level regulation and the broader gene-level enrichment within various pathways encompassing the innate immune response pathway and membrane trafficking.

### Novel transcript isoforms of familial AD genes and risk genes

The use of genome-wide long-read RNA-seq enabled the identification of several transcript variants originating from the canonical dementia causative and risk genes. Notably, novel transcripts of *Apoe, App*, Microtubule Associated Protein Tau (*Mapt*), and 2’-5’ Oligoadenylate Synthetase 1A (*Oas1a*) were discovered, that were previously absent in the Ensembl catalogue (Table S9). Among these was an anti-sense transcript represented by StringTie identifier MSTRG.40715.1 of *Apoe* (Figure 4, as compared to the reference annotation). While the various transcripts of these dementia causative and risk genes likely play distinct roles under specific conditions (e.g. during development), the changes in the levels of these isoforms at 9-months of age were not dependent on amyloid-β.

**Figure 4.**
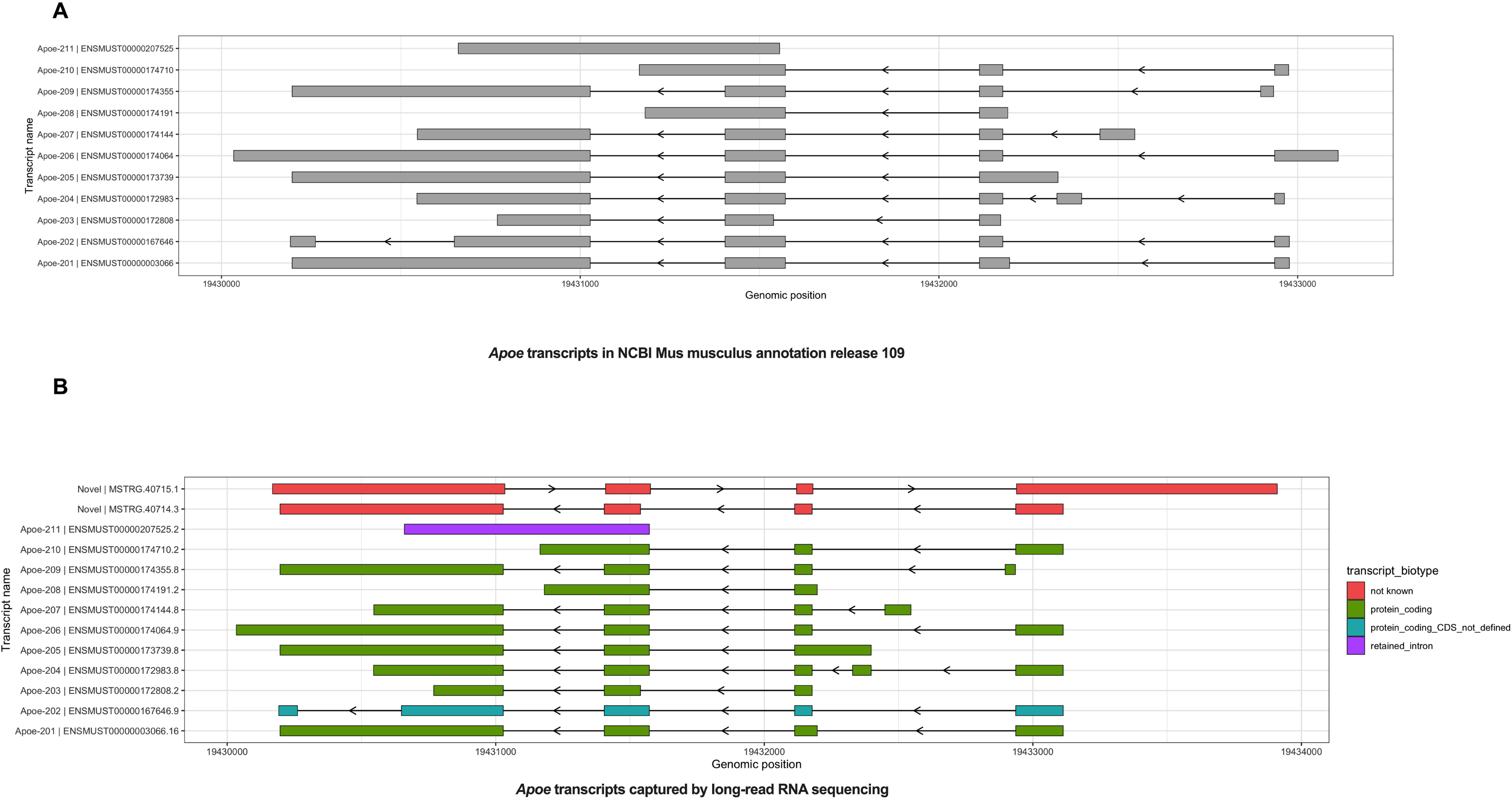
Integrative Genomic Viewer visualization of all detected *Apoe* transcripts. (A) Canonical *Apoe* transcripts in reference annotation (NCBI Mus Musculus 109). The common *Apoe-201* isoform is shown at bottom. (B) The isoforms detected via long-read RNA-seq represents two novel transcript variants in the *Apoe* gene. In addition to the canonical transcripts, our analysis uncovered two novel transcripts (MSTRG.40715.1, MSTRG.40714.3), including an antisense transcript aligned in the reverse direction of the reference genome.

### Detecting genome-wide splicing patterns, isoform switching, and isoform/exon usage using long-reads

We employed the DEXSeq analysis approach to investigate the intricate mechanisms underlying gene and transcript expression in *App^NL-G-F^* mice compared to controls. This method utilizes a negative binomial distribution to model the feature counts of each gene, such as exons, and incorporates Generalized Linear Models with interaction terms to capture the interplay between different features within the same gene^45,49^ (Table S10). To gain further insights into the landscape of alternative splicing, we initially performed differential transcript usage (DTU) and differential exon usage (DEU) analyses; which are calculations based on isoform and exon fraction per sample. Changes in isoform usage provide additional clarity on instances where the proportional contribution of isoforms (also known as isoform switching) relative to the overall gene expression undergo significant alterations between different genotypes. Isoform switch detection, and coding potential analysis, protein domain identification and external annotations of the novel and known isoforms have been performed via the IsoformSwitchAnalyzer^47^ tool. To further elucidate the functional ramifications of altered transcript/exon use and alternative splicing, and their potential effects on changes at the protein level, we employed the tappAS tool which is a part of the Functional Iso-transcriptomics Framework^48^, also accounting for differential polyadenylation usage of transcripts. In addition, we identified the genome-wide splicing profile using PSI-Sigma, a splicing detection tool specifically developed for long-read RNA-seq^46^. Collectively, the combination of genome-wide gene and isoform level analyses with more complete transcript detection and quantification enabled us to validate and identify alternative splicing events while providing predictions of their functional consequences. Notably, in addition to the preferential usage of specific transcript isoforms, our findings suggest that the selection of certain exons may play a critical role in refining mRNA regulation, cellular trafficking and potentially influencing protein function by changing protein sequence.

### Selective transcript usage and exon usage

Our genome-wide transcript isoform usage analyses using DEXSeq and tappAS identified a number of transcript isoform changes in response to amyloid plaques (Figure 5A, Tables S11 and S12), including several genes associated with the ARM/DAM phenotype, such as *Apoe* (ENSMUST00000174064.8, FDR= 2.59e^-03^), Colony Stimulating Factor 1 (*Csf1*, ENSMUST00000120243.8, FDR = 4.22e^-03^), Synaptogyrin 1 (*Syngr1*, ENSMUST00000009728.13, FDR = 6.17e^-34^), Colony Stimulating Factor 2 Receptor Subunit Alpha (*Csf2ra*, ENSMUST00000235172, FDR = 4.32e^-04^), Cell Adhesion Molecule 2 (*Cadm2*, ENSMUST00000114548, FDR = 7.59e^−03^), and Insulin-like growth factor 1 (*Igf1*, ENSMUST00000121952, FDR = 5.20e^-05^). Strikingly, the GO annotations associated with these genes exhibiting transcript preference were predominantly enriched in postsynapse organization, microtubule cytoskeleton, regulation of MAPK cascade, interferon-γ production, and intracellular signal transduction; shedding light on their role on synaptic pathways in response to amyloid pathology (Figure 5B). Furthermore, our analysis discovered multiple instances of exon usage differences in key genes such as *Ctsd*, Cathepsin B (*Ctsb)*, *Syngr1*, BCL2 Related Protein A1 (*Bcl2a1*), *App*, and Clusterin (*Clu*) in response to amyloid (Table S10). Particularly intriguing is the alternative splicing of *Clu*, which is known to modulate β-amyloid metabolism and/or deposition^50^.

**Figure 5.**
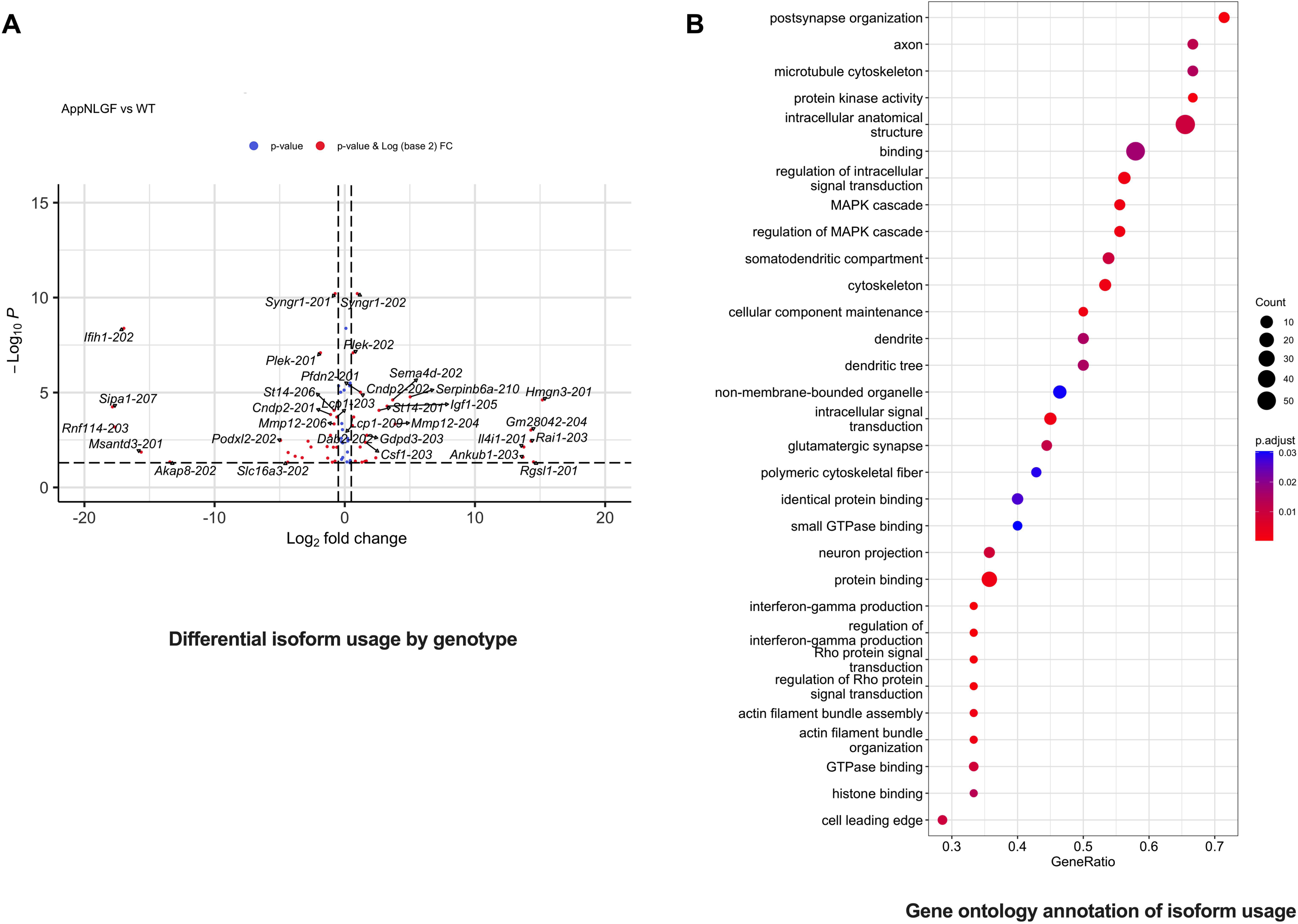
Differential transcript usage analysis of genes in *App^NL-G-F^* vs control mice at 9 months of age using long-read RNA-seq. (A) Analysis of transcript-level expression identified a total of 70 transcripts that exhibit preferential usage of specific isoforms in response to amyloid plaques. (B) Differentially transcript usage isoforms are enriched in postsynaptic organisation, regulation of intracellular signal transduction and MAPK cascade. (C) The GO enrichment analysis of the differential transcript usage list revealed associations with cell developmental pathways, negative regulation of amyloid fibril formation, and neuron projection extension. These findings diverge notably from the outcomes observed in both gene-level and isoform-level expression analyses.

Isoform switch results displayed a novel transcript variant in Capping Actin Protein (*Capg)* in *App^NL-G-F^* animals (denoted MSTRG.38755.8; Figures 6A and 6B), with decreased usage of *Capg* isoform ENSMUST00000071044.13, and increased usage of ENSMUST000000114071.8 compared to wildtype controls. The isoform switch between these three isoforms may exert functional consequences, as MSTRG.38755.8 is a coding isoform, that lacks a protein domain between positions 31-107 of the amino acid sequence (Figure 6C), which may regulate the functional role of CAPG. Several other genes including amyloid-responsive microglial genes, *Gusb* (glucuronidase beta; Figure 7A) and PTPRF Interacting Protein Alpha 4 (*Ppfia4*; Figure 7B) showed significant isoform switch events, including usage of the novel variant MSTRG.2151.1 of *Ppfia4*. We also detected that *Capg,* Transmembrane Protein With EGF Like And Two Follistatin Like Domains 1 (*Tmeff1*) and Occludin (*Ocln*) underwent switching of their polyadenylation sites in response to amyloid plaques (Figure 7C). Biological annotation of genes showing isoform switching indicated enrichment of pathways involving development, cytoskeletal protein binding, and cell junction (Figure S4).

**Figure 6.**
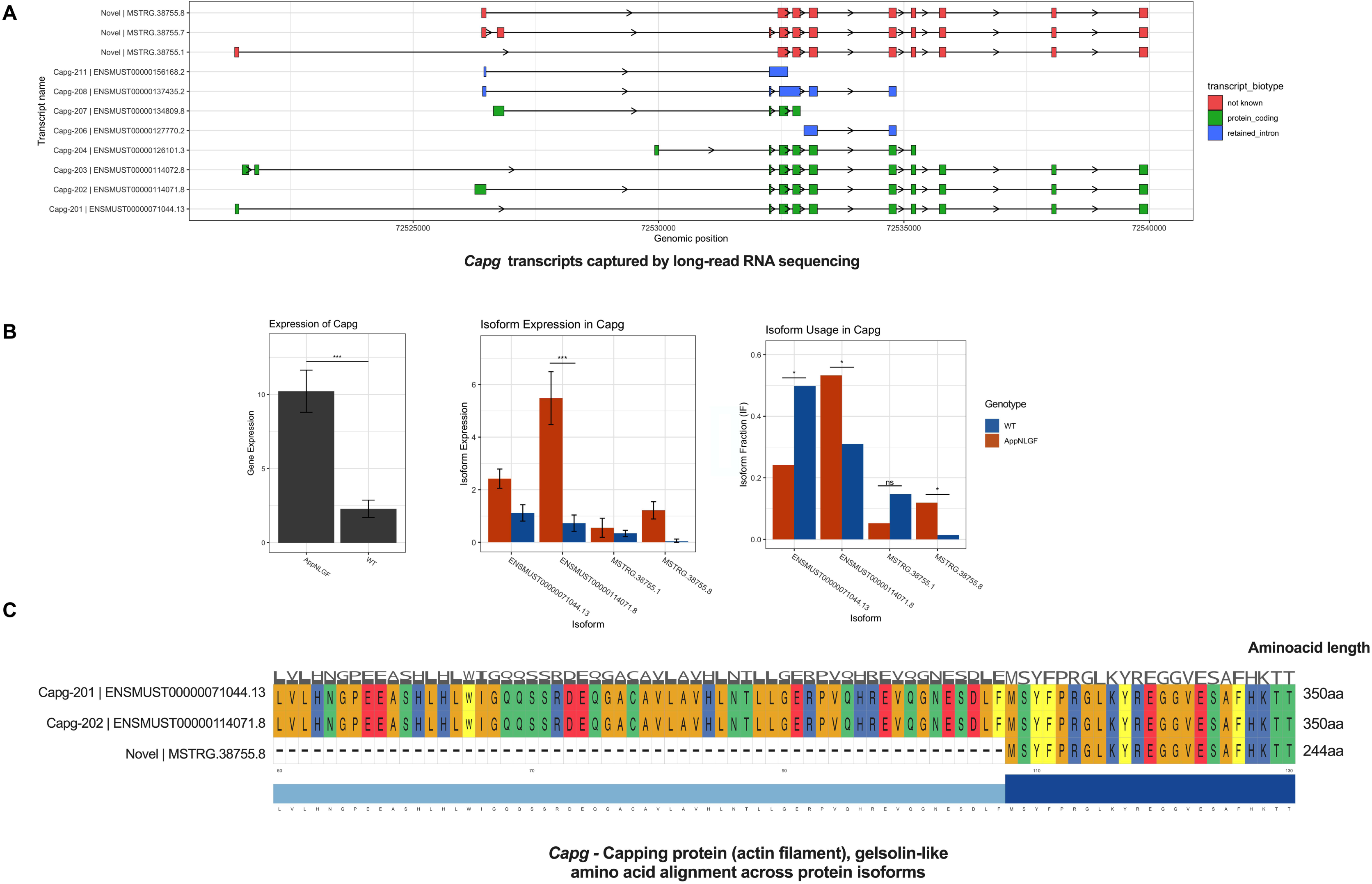
*Capg* showed isoform switching and amino acid changes in the predicted protein sequence in its novel transcript isoform. (A) The isoforms detected in the *Capg* gene. In addition to the canonical transcripts and common *Capg-201* form, three novel transcripts (MSTRG.38755.8, MSTRG.38755.7, MSTRG.38755.1) were found. (B) Gene-level expression of *Capg* is increased with amyloid pathology (left panel), while isoform expression specifically indicates individual isoforms that are upregulated in response to amyloid pathology (middle panel). Isoform usage analysis revealed an isoform-switch pattern in proportional transcript use between ENSMUST00000071004.13, ENSMUST00000114071.8 and novel transcript variant MSTRG.38755.8 (right panel). (C) The novel coding transcript (MSTRG.38755.8) lacks a full domain between 31-107 amino acids (https://www.uniprot.org/uniprotkb/Q3TNN6).

**Figure 7.**
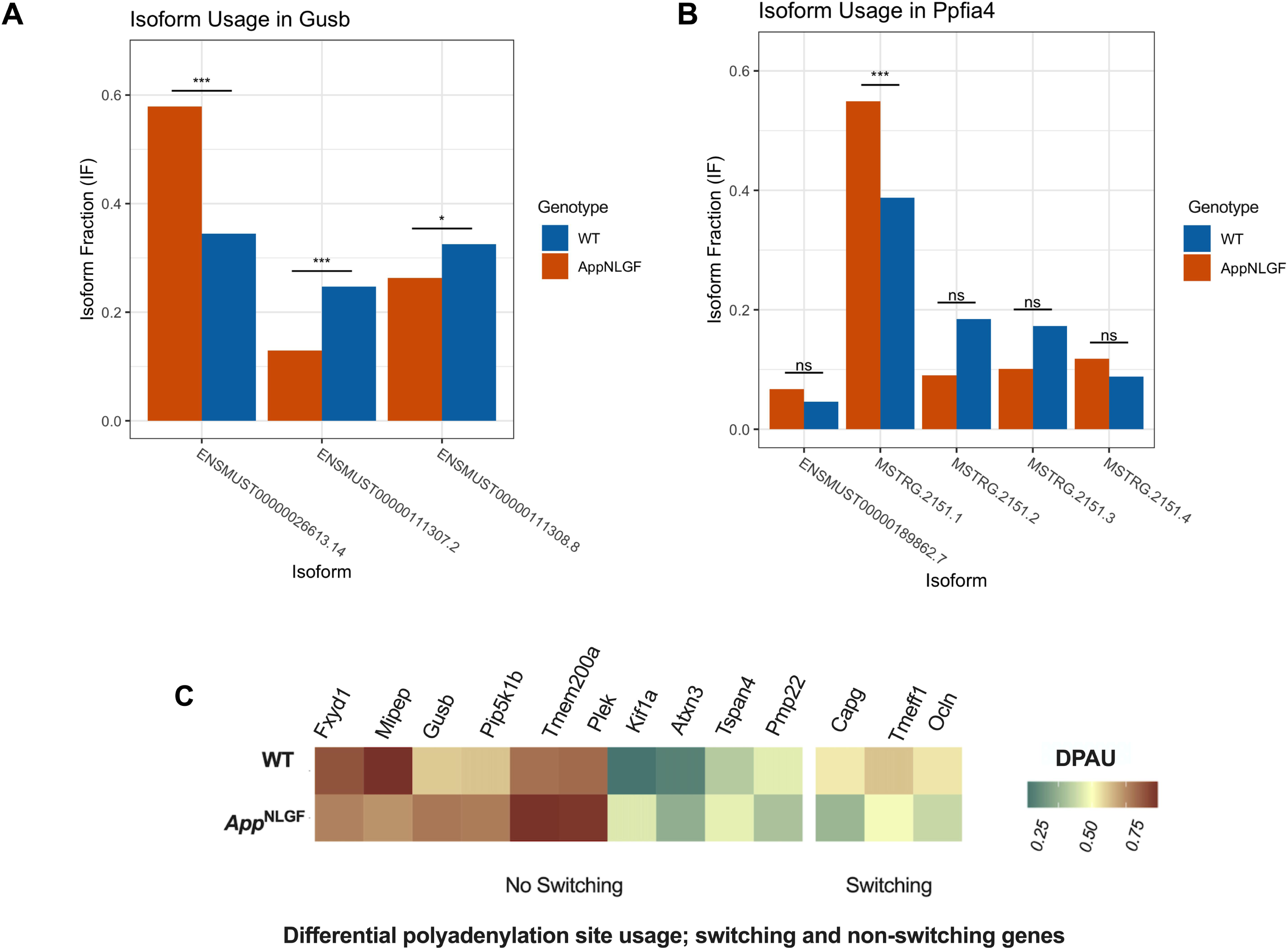
ARM/DAM gene *Gusb* and trans-synaptic signalling gene, *Ppfia4,* were amongst alternatively spliced genes showing transcript usage and isoform switching. (A) *Gusb* showed an isoform switch event in proportional transcript usage (FDR = 2.5e^-06^), a single exon skipping event on Exon 3 of ENSMUST00000026613.14 (FDR = 1.0e^-03^) and changes in exon usage (FDR = 4.1e^-02^) in *App^NL-G-F^* mice. (B) *Ppfia4* showed an isoform switch event for MSTRG.2151.1 (FDR = 3.7e^-07^), and an exon skipping event on Exon 3 (FDR = 5.9e^-04^) which potentially increases overall isoform expression in response to amyloid. (C) Altered polyadenylation sites in genes, including *Capg, Tmeff*, and *Ocln*, demonstrated a switch in polyadenylation patterns between *App^NL-G-F^* and control samples. This observed change in differential polyadenylation site usage (DPAU) is suggestive of alterations in mRNA stability, subcellular localization, RNA-protein binding, and/or translation efficiency, indicating potential regulatory modifications at the post-transcriptional level^109–111^.

There are numerous usage changes of specific exons of the abovementioned genes that lead to either changes in isoform usage, and/or isoform switching (Tables S11, S12 and S13). Changes in exon usage have the potential to lead to specific isoform preferences, as exemplified by the *Syngr1* gene. We observed an increased usage of Exon 7, Exon 9, and Exon 10 (Stage-wise testing^51^, FDR < 0.05), concomitant with decreased usage of *Syngr1-201,* and increased usage of *Syngr1-202* isoforms in response to amyloid. This isoform switch (FDR = 1.21e^-37^) highlights the dynamic nature of alternative splicing and suggests a potential functional implication in the context of amyloid pathology.

Cytoplasmic FMR1 interacting protein 2 (*CYFIP2*) demonstrated decreased protein expression and is associated with memory loss in the advanced stages of human AD^52^. In our study, we observed alterations in a mouse model representing the early stages of AD, specifically, we noted changes in the usage of *Cyfip2-205* (Stage-wise testing^51^, FDR = 1.82e^-05^). This mechanistic insight might signify an early phase response to amyloid marked by preferential expression of specific mRNA isoforms, eventually culminating in the accumulation of tau tangles and transition to the later stages of the disease with neurodegeneration^53–55^.

### Alternative splicing analysis in response to amyloid plaques serves to adapt synaptic function

To identify different alternative splicing events more directly, we used the PSI-Sigma software to classify splicing events into five standard types: single or multi-skipped exons (SES/MES, also known as cassette exons), alternative 5’ and 3’ splice sites (A5SS and A3SS), intron retention (IR), and mutually exclusive exons (MXE)^46^. We identified approximately one hundred genes displaying one of these classes of alternative splicing in response to amyloid (Table S14). Interestingly, we identified single/multi-exon-skipping events in several ARM/DAM genes, including *Ctsd*, Cathepsin A (*Ctsa*), Glycosylated Type I Transmembrane Glycoprotein (*Cd68*), Glucuronidase Beta (*Gusb*), Tumor Protein D52 (*Tpd52*), and Cytokine Receptor Like Factor 2 (*Crlf2*). Additionally, we found that *Csf2ra* and LYN proto-oncogene, Src family tyrosine kinase (*Lyn*) exhibited alternative 5’ UTR splicing sites with amyloid pathology. Overall splicing increased with amyloid pathology, with a mean change in Percent Spliced-In (ΔPSI) of 1.76% for all significant splicing events (p < 0.01 and |ΔPSI| >= 5%) between *App*^NL-G-F^ and wildtype samples. The genes exhibiting alternative splicing in response to amyloid were found to be enriched for functions primarily associated with endomembrane system organisation, actin filament binding, actin cytoskeleton reorganisations and kinase binding (Figure S4).

To validate the isoform usage and splicing changes we identified, we mined short-read RNA-seq data across three prominent human AD cohorts: Mayo Clinic, Mount Sinai Brain Bank (MSBB), and the Religious Orders Study (ROS) and Rush Memory and Aging Project (MAP) (ROSMAP)^30^. Marques-Coelho and colleagues used a combination of differential gene expression and isoform switch analyses to identify differential transcript usage from short-read RNA-seq of temporal and frontal lobes in healthy and AD adult individuals^28^. Although the conclusions made from short-read RNA-seq can be limited, where information obtained is short range from adjacent exons, we found that a number of genes showing isoform usage changes in mice in response to amyloid *also* showed isoform usage changes in late-stage AD brain, including genes such as *Ctsa,* Clathrin Light Chain A (*Clta*), DENN Domain Containing 2A (*Dennd2a*), Interferon Regulatory Factor 9 (*Irf9*) and SMAD Family Member 4 (*Smad4*)^30^.

### Transcript variation and splicing reveals changes to numerous cell-types including synaptic changes during early amyloid accumulation

Our detailed analyses on transcript usage, exon usage and alternative splicing offer novel insights into the functional implications of transcript isoform switching during microglial activation, synaptic adaptation and a series of other cell type changes in response to amyloid plaques in models of early AD. Importantly, these transcript isoform modifications exhibit widespread enrichment across various cell types present in the brain as illustrated in the cell-type enrichment heatmap in Figure 8. This includes astrocytes, oligodendrocytes, both excitatory and inhibitory neurons, and T cells^56–59^. A particularly intriguing aspect is the enrichement of genes with isoform-level changes associated with diverse synaptic pathways. This suggests concurrent alterations in synaptic physiology alongside the upregulation of AD risk genes in microglial proliferation and activation, during the initial stages of disease progression, aligning with the earliest cognitive changes. These findings hold the potential to provide more precise isoform-specific indicators of disease stage for diagnostic tracking purposes, independent of the gene-level expression changes resulting from microglial cell proliferation/activation.

**Figure 8.**
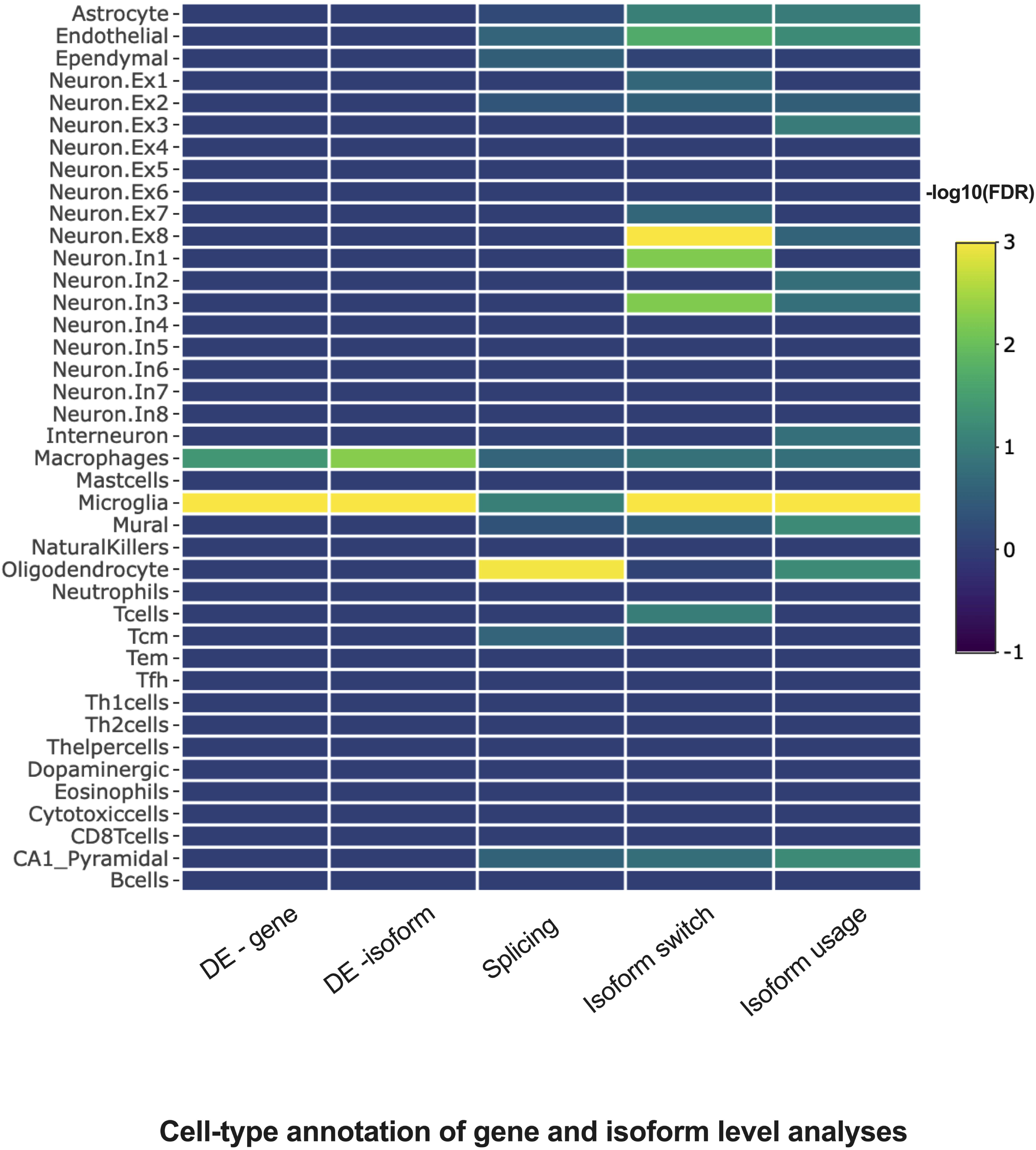
Heatmap of cell-type enrichment of gene and isoform level analyses. Gene-level differential expression (DE) analysis reveals an enrichment of genes and isoforms within those expressed by microglia and macrophages, primarily attributed to the heightened proliferation and activation in response to amyloid accumulation. Genes showing splicing events were predominantly expressed by oligodendrocytes. Isoform switching mainly occurs in genes expressed by neuronal subtypes, endothelial cells, mural cells, and astrocytes. Isoform usage changes occur in genes expressed by interneurons, mural cells, neuron subtypes, endothelial cells, and astrocytes^57,58^. Overall, this figure underscores the intricate profile of isoform-level changes across different cellular environments, a complexity that is not fully captured by gene- or isoform-level differential expression analyses alone.

## Discussion

### Microglial activation, AD risk genes and alternative mechanisms linking to synapses

GWAS have transformed how we view AD, which has led to a shift in focus towards the investigation of the innate immune system. These studies have identified around 100 genes/loci variants associated with AD risk^2–5^, many of which are expressed by microglia in the brain, implicating the innate immune system in AD pathogenesis^4^. The genetic network formed by these risk genes is induced by amyloid pathology, and in large due to triggering of microglial proliferation and activation^16^. Intriguingly, this microglial risk network activation occurs early in disease progression, preceding neuronal death and cognitive decline^60^. Exploring how these risk genes/loci influence AD susceptibility includes exploring alternative mechanisms when variants do not alter the protein coding sequence of genes. Non-coding variants, such as expression quantitative loci (eQTLs) and splicing quantitative loci (sQTLs), can modulate gene expression and splicing patterns. This can influence transcript abundance, specific transcript isoforms, subcellular localisation of RNA, and changes in the proteome to compound AD risk^61,50^. While short-read RNA-seq is valuable for quantifying gene-level expression changes, it has limitations in capturing differential transcript usage, exon usage and alternative splicing. To address such limitations, we performed long-read RNA-seq of cortical RNA in the *App^NL-G-F^*mouse model of amyloid plaque accumulation compared with wildtype control samples. Analysis of gene expression was undertaken at 9-months of age in mice that have already developed a modest amyloid-β associated spatial short-term memory deficit. We identified novel transcripts from causal/familial and risk genes associated with AD. We observed induction of microglial-expressed AD risk genes at the whole gene-level, but also discovered preferential transcript usage in microglial genes in response to amyloid plaques. Moreover, a significant number of the genes which exhibited differential exon usage and alternative splicing in response to amyloid, were enriched in functional pathways coordinating synaptic physiology and the interaction with diverse cell types in the brain encompassing T cells, astrocytes and oligodendrocytes. Understanding these novel changes at the transcript-level will provide new insights into microglial activation, synaptic adaptation and interaction between other cell types at early disease stages, potentially informing disease-stage and isoform-specific diagnostic and therapeutic strategies.

### Early spatial short-time memory and retinal deficits in the *App^NL-G-F^* knock-in mouse model

We found a progressive spatial short-term memory deficit in the *App^NL-G-F^*knock-in mouse model of amyloid-β deposition. This is consistent with previous reports of deficits in place preference memory^62^ and spatial memory as tested in the Barnes maze^63^ at 8-months of age. In contrast, we did not find deficits in short-term novelty recognition memory at the same age, consistent with a previous report^64^ that did not find deficits in long-term (24 hours) recognition memory in male *App^NL-G-F^* mice at 6-months of age. Notably under some testing paradigms (1-hour inter-trial-interval) short-term recognition memory has been reported to be impaired in this model at 10-months of age^65^. Differences in the testing paradigm used in this study, previous experience of the mice and differences in husbandry conditions may have contributed to these differences. Overall, comparative analysis of the spatial novelty versus novel object recognition revealed a specific spatial short-term memory deficit in *App^NL-G-F^* model of AD. This may relate to vulnerability of cellular populations in the hippocampus and dorsolateral prefrontal cortex to pathogenic amyloid-β in *App^NL-G-F^* mice, as important brain areas for spatial memory storage and maintenance^66^, which is in good agreement with clinical findings on AD patients^67^. In parallel, we observed subtle changes to retinal structure of the *App^NL-G-F^* knock-in mouse model, with a thinner ganglion cell layer in female mice. This is inconsistent with a previous study^68^ of female mice that found no significant differences in retinal structure or alterations to visual function in the *App^NL-G-F^* model. In older mice amyloid-β has been reported to accumulate in the retina of this model^68,69^. Further research is thus warranted to verify the changes to the retinal structure we observed and to determine if these impact visual function and behavioural outcomes in this model. In summary, the *App^NL-G-F^* mice at 9 months of age showed early AD-relevant behavioral changes, thus the observed changes in alterative splicing and isoform usage observed may have functional consequences.

### Differentially expressed genes and transcripts in long-read RNA-seq bulk experiments are mostly microglial

Our gene-level expression analysis, employing long-read RNA-sequencing, corroborated short-read data from other mouse models, highlighting induction of *Trem2*, Tyro protein tyrosine kinase-binding protein (*Tyrobp*), and other genes associated with the ARM/DAM microglial cell-state and AD risk. The utility of long-read RNA-seq in capturing gene-level expression changes underscored microglial proliferation and activation in this mouse model of amyloid accumuation. Importantly, our behavioural analysis demonstrated that these gene-level expression changes coincide with early cognitive alterations, suggesting their relevance in disease progression from early stages. Notably, several ARM/DAM genes were found to be upregulated, further supporting the involvement of microglia in the response to amyloid pathology.

While gene-level changes provide insights into the amyloid-responsive genetic network expressed by microglia, our analysis reveals that specific microglial genes exhibit preferential selection of transcript isoforms in response to amyloid. Remarkably, these genes demonstrate notable disparities in fold-expression ranking and significance when compared at the gene-level versus the transcript level, as exemplified by genes such as *Igf1*. The differential usage of specific transcripts by these genes in the presence of amyloid plaques suggests their potential role in driving the cellular response to amyloid. It is worth noting that *Igf1*, one of the genes showing selective transcript usage, plays a crucial role as a mediator in the clearance and regulation of amyloid-β in the brain, highlighting its relevance in the context of AD progression^70^.

A further important example is the distinction between the mouse *Trem2* isoforms: we detected three isoforms of *Trem2*, the ENSMUST00000024791.14 isoform, encoding for the full-length receptor, and the ENSMUST00000113237.3 isoform, lacking the transmembrane-receptor, which are both induced by amyloid plaques in our analysis. In contrast, the ENSMUST00000132340.2 isoform which represents an intron-retention event is not thought to produce protein and is not induced by amyloid. Focussing on specific isoforms is functionally important, the human *TREM2* isoform ENST00000373113, responsible for encoding the full-length transmembrane receptor, along with the alternatively spliced isoforms ENST00000373122 and ENST00000338469, demonstrate a moderate increase in specific brain regions among individuals with AD^71^. Alongside this, experimental findings from 7-month-old control mice and 5xFAD mice, amyloid-ß^71^ expressing the human *TREM2* gene, compared to control mice (B6^hT2^ and 5xFAD^hT2^), provide evidence that the alternatively spliced isoforms of *TREM2* undergo translation and secretion, leading to the formation of soluble TREM2 (sTREM2).

Beyond microglia, the most abundantly expressed *Gfap* isoform detected in our dataset, was also found to be major driver of the *Gfap* expression in a transgenic mouse model of tau pathology after long-read RNA-seq analysis^72^, suggesting that the role of this isoform in the role of the astrocyte response in AD may in particular warrant further investigation. Thus identifying specific transcript isoforms provides new biological insights into mechanisms leading to AD.

### Splicing and preferential transcript usage reveal insights into synaptic adaptation

Accompanying the well-characterised induction of microglial-expressed AD risk genes at the gene-level in response to amyloid, we also revealed preferentially used novel isoforms, transcript usage, switching and splicing events in a series of microglial genes. These included many ARM/DAM cell-state genes associated with amyloid pathology, independent from changes in cell number due to proliferating microglia. Transcript isoform changes were seen in genes such as *Capg*, *Trem2*, *Ocln, Ctsd, Ctsa, Cd68, Gusb*, *Csf2ra* and *Ctsb*. Recent work has shown that changes in CTSB activity may contribute to AD^73^. Thus intricate control of transcript isoforms of these microglial genes within AD risk genetic pathways/networks is likely to tailor the functional microglial response to amyloid. To investigate if genes that exhibited transcript isoform alterations due to amyloid, as identified through our long-read sequencing in the *App^NL-G-F^* mouse model, also displayed similar changes in orthologous genes in human AD, we analysed short-read RNA-seq data from the Mayo Clinic, MSBB, and ROSMAP^30^. To identify differential transcript usage from temporal and frontal lobes in healthy and AD adult individuals^30^, Marques-Coelho and colleagues used isoform switch analysis. Mining this data, we found a number of genes showing isoform usage changes in late stage AD brain that also showed usage changes in mice in response to amyloid. By comparing short-read and long-read RNA-seq, as well as examining differences between human and mouse, and late-stage human AD and early amyloid pathology, we observed that at least part of the response to AD-related pathology was conserved. This response involves mechanisms related to transcript usage and splicing in shared genes, including *Ctsa, Clta, Dennd2a, Irf9 and Smad4*. Appreciation of transcript isoform control, and more accurate identification of the novel and “in-catalogue” isoforms with long-read analysis, will be important to understand the mechanisms by which microglia control AD risk.

We explored the splicing events and differentially used transcripts associated with amyloid pathology, extending beyond microglial proliferation due to amyloid. Our findings are supported by relevant studies shedding light on the functional significance of these genes in AD. For instance, *SEMA4D* (*CD100*) plays a regulatory role beyond axon guidance, and class 4D in particular is associated with immune modulation^74–76^. Its upregulation has been linked with neuronal stress and astrocyte responses^77^. Our analysis revealed the preferential upregulation of specific isoforms of *Sema4d*, along with concomitant alternative splicing events such as the expression of A5SS *(Sema4d-204)* and single-exon skipping. Additionally, other genes involved in synaptic physiology, such as *Syngr1* and *Cyfip2*, exhibited multiple differential transcript usage and exon usage changes in response to amyloid. Notably, *SYNGR1,* a synaptic vesicle-associated trafficking protein, found to be increased in synaptoneurosomes from the prefrontal cortex of AD patients compared to controls^78^. It is possibly involved in compensatory *APOE*-dependent synaptic plasticity^79^. *Syngr1* knockout mice show no overt changes in synaptic architecture, whilst electrophysiological analysis of these mice shows impaired hippocampal short- and long-term synaptic plasticity^80^. The the depletion of *Syngr1* along with other “gyrin and physin proteins” caused defects in synaptic plasticity, where deletion of all led to increased synaptic vesicle release^81^. Thus*, SYNGR1* is potentially involved in synaptic modifying activity-dependent synaptic transmission, and may play a crucial role in synaptic adaptation in response to amyloid. The Myeloid Landscape study demonstrated stronger expression of *Syngr1* in microglia compared to neurons in mice (Myeloid Landscape^82^), thus *Syngr1* pathology-associated isoforms may be acting through both synapses and microglia. Further research is warranted to determine if the observed changes in synaptic gene expression contribute to the observed changes in spatial learning observed in the *App^NL-G-F^* mouse model.

A further example of selective transcript usage is *Cyfip2*, a regulator of actin dynamics at synapses^83,84^, which is postulated to bridge amyloid and tau pathology. Our study highlighted differential usage of *Cyfip2-205* compared to other transcripts, further emphasizing its role in synaptic adaptation^78^ in response to early amyloid deposition in our model^44^. The observed interplay between splicing alterations alongside the upregulation of AD risk genes suggests a reciprocal relationship, possibly impacting synaptic responses to amyloid. This intricate interplay proposes microglial AD risk gene variants may modulate synaptic responses through the splicing events we identified via long-read sequencing. Our findings enhance understanding of the synaptic adaptation in AD and underscore the potential of splicing events and preferential transcript usage as diagnostic biomarkers and therapeutic targets.

### Implications for disease staging and therapeutic interventions

Discovering preferential transcript usage, alternative splicing, and novel transcripts in response to amyloid pathology may hold significant implications for AD diagnosis, disease staging, and therapeutic interventions. While we have not directly studied isoform changes over time, it is possible that these novel isoforms provide addtitional biomarker targets, and that monitoring alternative splicing, or isoform usage, over time could provide information related to disease stage, progression or response to intervention. It may be possible to target specific transcript isoforms linked to synaptic protection or dysfunction to inform new therapeutic interventions, such as antisense oligonucleotides, to modulate splicing patterns and switch between detrimental and protective isoforms^85^. Such interventions could hold promise for attenuating disease progression and improving synaptic function, opening up new avenues for therapeutic exploration in AD.

## Methods

### Animal welfare and husbandry

All animals were housed and maintained in the Mary Lyon Centre, MRC Harwell Institute, under specific pathogen-free (SPF) conditions, in individually ventilated cages adhering to environmental conditions as outlined in the Home Office Code of Practice. All animal studies were licensed by the Home Office under the Animals (Scientific Procedures) Act 1986 Amendment Regulations 2012 (SI 4 2012/3039), UK, and additionally approved by the Institutional Ethical Review Committees. All mice were co-housed throughout the study as lone housing is known to modify amyloid-β associated phenotypes^86^; mice were housed with animals of the same genotype and sex, weaned at the same time, to prevent phenocopying. Mice had access to a cardboard tunnel with bedding material and wood chips (grade 4 aspen). All animals had continual access to water and RM3 chow (Special Diet Services). Animals were euthanized by terminal perfusion with PBS (under sodium pentobarbitone (Euthatal) anaesthesia) in accordance with the Animals (Scientific Procedures) Act 1986 (United Kingdom).

### Animal genetics and experimental design

The *App*^tm3.1Tcs^ (MGI:5637817)^38^, herein referred to as the *App^NL-G-F^*mouse model was maintained on a C57BL/6J genetic background as a heterozygous backcross. Cohorts of homozygous and wildtype (WT) controls were generated by heterozygous cross for this study. The number of animals used in the phenotyping pipeline were: *App*^+/+^ (WT allele homozygotes) male n = 23, female n = 21; *App^NL-G-F/NL-G-F^* (mutant allele homozygotes) male n = 15, female n = 24. Subsets of these animals were used for long-read sequencing analysis (*App*^+/+^ male n = 3, female n = 3; *App^NL-G-F/NL-G-F^*male n = 3, female n = 3), and biochemical amyloid-β quantification (*App*^+/+^ - male n = 6, female n = 6, *App^NL-G-F/NL-G-F^* male n = 6, female n = 6).

### *In vivo* phenotyping study design

The cohort was longitudinally tested in the following order: ECHO-MRI (8 weeks of age); Home Cage Analysis (10 weeks of age); Forced Y-maze (12 weeks of age); Novel Object recognition (14 weeks of age); Optical Coherence Tomography (15-16 weeks of age); Auditory Brainstem response (15-16 weeks of age); ECHO-MRI (17 weeks of age); Home Cage Analysis (32 weeks of age); Forced Y-maze (33 weeks of age); Novel Object recognition (34-35 weeks of age); Optical Coherence Tomography (37-38 weeks of age); ECHO-MRI (39 weeks of age). All behavioural testing was undertaken between 08.00 -16.00 (light 7.00–19.00 and dark 19.00–7.00). All phenotyping and data analysis was performed blind to genotype.

### Auditory brainstem response (ABR)

Auditory brainstem response (ABR) tests were performed using a click stimulus in addition to frequency-specific tone-burst stimuli at 6, 12, 18, 24 and 30 kHz, as described previously^41^. Briefly, mice were anaesthetized by intraperitoneal injection of ketamine (100 mg/ml at 10% v/v) and xylazine (20 mg/ml at 5% v/v). Once anaesthetized, mice were placed on a heated mat inside a sound-attenuated chamber (ETS Lindgren) and recording electrodes (Grass Telefactor F-E2-12) were placed subdermally over the vertex (active), right mastoid (reference) and left mastoid (ground). ABR responses were collected, using TDT System 3 hardware and BioSig software (Tucker Davies Technology, Alachua, FL, USA). Stimuli were presented to the right ear of the anesthetized mouse. Auditory thresholds were defined as the lowest dB SPL that produced a reproducible ABR trace pattern and were determined visually.

### Echo Magnetic resonance imaging (ECHO-MRI)

Body composition analysis of mice was determined using an EchoMRI whole body composition analyser (Echo Medical System, USA). Fat mass and lean mass were recorded as well as weight measured. However subsequent analysis indicated that the presence of the RFID chip used in Home Cage Analysis procedure (see below) interfered with the fat mass recording.

### Spatial novelty preference in the forced Y-maze

Spatial novelty preference was assessed in an enclosed Perspex® Y-maze as described previously^87^. Briefly, a Perspex® Y-maze with arms of 30 × 8 × 20 cm was placed into a room containing a variety of extra-maze visual cues. Mice were assigned two arms (the ‘start’ and the ‘other’ arm) to which they were exposed during the first phase (the exposure phase), for 5 minutes. Timing of the 5 minutes period began only once the mouse had left the start arm. The mouse was then removed from the maze and returned to its home cage for a 1 minute interval between the exposure and test phases. During the test phase, mice were allowed free access to all three arms. Mice were placed at the end of the start arm and allowed to explore all three arms for 2 minutes beginning once they had left the start arm. An entry into an arm was defined by a mouse placing all four paws inside the arm. Similarly, a mouse was considered to have left an arm if all four paws were placed outside the arm. The amount of time spent in each arm, the number of entries made into each arm and the total distance moved were recorded using EthovisionXT 8.5 (Noldus). A novelty preference ratio was calculated for the time spent and number of entries in arms [novel arm / other arm)].

### Novel Object Recognition

Recognition memory was assessed using novel object recognition. Three consecutive days prior to testing mice were habituated to the test arena by allowing each mouse to freely explore the arena for 10 minutes. The test arena was an enclosed Perspex® opaque arena measuring 44 x 44 cm. Habituation and testing was performed at a light level of 20 lux, to minimise anxiety. On the day of testing the arena was set up with two identical objects, each placed 10 cm away from two adjacent corners. The mouse was allowed to explore the arena and objects for 10 minutes (exposure phase). The number and length of investigations made by the animal to each object was scored using EthovisionXT 8.5 (Noldus). Analysis of this data found no significant genotype differences in object exploration during the exposure phase. The animal was then returned to the home cage for 2 minutes. Following this, the arena was set up with one familiar object (a new copy of the object used in the exposure phase) and one novel object, each placed 10 cm away from two adjacent corners. The mouse was then allowed to explore the arena and objects for 5 minutes (test phase). During this time the number and length of investigations made by the animal to each object was recorded using EthovisionXT 8.5, and approaches were manually scored by an investigator blinded to sex and genotype. The test objects used were a glass bottle and wood block for testing at 14 weeks and plastic bottle and tin can for testing at 34-35 weeks. Object assignment and location within the arena was randomly assigned to each animal.

### Optical Coherence Tomography

Retinal morphology was assessed by Optical Coherence Tomography (OCT) using a Leica Bioptigen system according to manufacturer’s instructions. Briefly, animals were anaesthetized using 10µl/g ketamine/xylazine solution subcutaneously in the scruff and a two-dimensional image of the retina was acquired using the Leica Bioptigen system. Tropicamide was used to dilate the pupils for image acquisition on both eyes. The mice were then positioned on the restraining cassette, the front teeth of the mice were placed on the bite bar. The cassette was then positioned in front of the lens and a drop of lubricant was applied to both eyes. The animals were then positioned so that the image was centred to the optical nerve head prior to beginning image acquisition. Images were acquired in Envisu R2200 Bioptigen System software, using a 1.4x1.4mm rectangular perimeter with 100 frames/B-scan. Following data capture, retinal layer segmentation was performed using OCTExplorer 3.8.0 (https://www.iibi.uiowa.edu/oct-reference).

### Microchipping

Radio frequency identification microchips were injected subcutaneously into the lower left or right quadrant of the abdomen of each mouse at 7 weeks of age. These microchips were contained in standard ISO-biocompatible glass capsules (12 x 2.12mm; ID K162 transponder, AEG Identifikationssysteme GmbH). The procedure was performed on isoflurane sedated mice (Isoflo; Abbott, UK) after topical application of local anesthetic cream on the injection site prior to the procedure (EMLA Cream 5%; AstraZeneca, UK) and the entry wound sealed with a topical skin adhesive, (GLUture; Zoetis) at the end of the procedure. All injections were performed in a surgical suite or suitably aseptic work space. The animals were allowed to recover from the microchip procedure for at least one week before being placed in the HCA rigs for data collection. The procedure has been described previously in Hobson *et al* (2020)^88^.

### Home cage Assessment

Group housed animals were monitored as described in Bains *et al.* 2016^40^. Briefly group housed mice (three per cage) were tagged with RFID microchips and placed in the Home Cage Analyzer system (Actual Analytics, Edinburgh) for 72 hours, which captured movement of mice by both video tracking and tracking of RFID chips. Data collection was started at first lights off on day 0 and stopped at 1 hour after final lights on, day 3.

### DNA extraction and genotyping

DNA was extracted from ear biopsy, isolated at P14 using TaqMan Sample-to-SNP (Applied Biosystems). Mice were genotyped for the *App^NL-G-F^*allele using TaqMan multiplexed qPCR for the NL mutant and WT *App* alleles and *Dot1l* reference allele; using the following primers (forward 5′-GGAAGAGATCTCGGAAGTGAAGA-3′; reverse 5′-CAGTTTTTGATGGCGGACTTCAA-3′) and probes (5′-FAM-TGGATGCAGAATTCGGACATG-BHQ1-3′) for the *App* WT allele and the following primers (forward 5′-CGGAAGAGATCTCGGAAGTGAATCT-3′; reverse 5′-ACCAGTTTTTGATGGTGGACTTCA-3′) and probes (5′-FAM-AGATGCAGAATTCAGACATGATTC-BHQ1-3′) for the *App^NL^* mutant allele and *Dot1l* allele primers (forward 5′-TAGTTGGCATCCTTATGCTTCATC-3′; reverse 5′-GCCCCAGCACGACCATT-3′) and probe (5′-VIC-CCAGCTCTCAAGTCG-MGBNFQ-3′). G and F mutant and WT *App* alleles were genotyped using allelic discrimination assays; using the following primers for the *App* WT/G allele (forward 5′-CGATGATGGCGCCTTTGTTC-3′; reverse 5′-GTTGCCTCTTGCGCTTACAG-3′) and probes (*App*-WT allele 5′-TET-ACCCACATCTTCAGCAA-BHQ1-3′ and *App*-mutant (G) allele 5′-FAM-CCACATCTCCAGCAAA-BHQ1-3′) and the following primers for the *App* WT/F allele (forward 5′-GTGGGCGGCGTTGTCA-3′; reverse 5′-CGCCATGATGGATGGATGTGTA -3′) and probes (*App*-WT allele 5′-FAM-AGCAACCGTGATTGTCAT-BHQ1-3′ and *App*-mutant (F) allele 5′-TET-AGCAACCGTGTTTGTC-BHQ1-3′)^90^.

### Tissue preparation

Following perfusion with PBS (pH 7.4) brain samples were dissected and divided along the sagittal midline with one hemisphere immersion fixed in 10% NBF (neutral, phosphate buffered formalin) for a minimum of 48 hours prior to tissue processing and paraffin embedding. Once dissected, tissue samples were dehydrated, cleared, and processed into paraffin wax using a Tissue-Tek VIP 6 AI (Sakura) and finally embedded into paraffin wax blocks using a HistoCore Arcadia (Leica). The remaining hemisphere was further dissected into the hippocampal and cortex regions which were then placed into cryotubes and immediately snap frozen in liquid nitrogen.

### Immunohistochemistry of mouse brain

Wax embedded (FFPE) blocks were trimmed dorsally to give a coronal section of either the ventral or dorsal hippocampus. For each FFPE block, two 4 μm tissue sections (40 μm apart) were mounted onto SuperFrost Plus slides for subsequent staining and analysis. First, sections were deparaffinized and rehydrated in a Gemini AS Automated Stainer (Epredia) using xylene and a series of ethanol baths (100%, 95%, 85%) and washed with distilled water. Sections were then pretreated with 80% formic acid for 8 minutes. After pretreatment, a Ventana ULTRA automated stainer was used for the following; heat-induced epitope retrieval for 16 minutes at 100°C in Tris boric acid EDTA buffer (Ventana Medical Systems, 06414575001); endogenous peroxidases were quenched using Inhibitor D (DABMap™, Ventana Medical System); primary antibody incubation was preformed using biotinylated mouse monoclonal IgG1 antibodies against amyloid-β (82E1, IBL, 0.2 μg/ml) for 8 hours at room temperature; chromogen visualization was achieved using Ventana DABMap™ kit (Ventana Medical Systems); counterstained with Haematoxylin II (4 minutes, Ventana Medical Systems ) and bluing agent (4 minutes, Ventana Medical Systems). Stained slides were washed with a mild detergent and dehydrated/cleared in a Gemini AS Automated Stainer using a series of ethanol baths (85%, 95%, 100%) and xylene. Stained slides were then mounted onto coverslips using a ClearVue Coverslipper (Epredia) before slide scanning with Zeiss Axio Scan Z1 slide scanner.

Regions of interest (ROI) of cortex and hippocampus for each tissue section were selected from the scans using QuPath^89^. Analysis of ROI images was conducted with an ImageJ (RRID:SCR_003070)^90^ macro. Briefly, a standard threshold of pixel value was used to create a binarized mask (Binary Mask plugin) of all ROI images. Pixels above the threshold were identified as Aβ staining, while pixels below the threshold were identified as not containing Aβ staining. ImageJ (RRID:SCR_003070) quantified the percentage of the pixels within the ROI covered by the mask (Analyze Particles plugin), thus positive for staining.

### Tissue fractionation for amyloid-β MSD assay

Total cortical proteins were fractionated based on the method in Shankar *et al.* (2009)^91^. Total cortex was homogenized in three volumes of the weight of the heaviest sample of ice-cold Tris-buffered saline (TBS) (50 mM Tris-HCl pH 8.0) plus complete protease and phosphatase inhibitors (Roche). Homogenates were centrifuged at 186 000 *g* at 4°C for 30 min, and the resultant supernatant (the soluble TBS fraction) was stored at −80°C. The resultant pellet was homogenized in three volumes times the heaviest sample of 1% Triton™ X-100 in TBS plus complete protease and phosphatase inhibitors (Roche) and centrifuged at 186 000 *g* at 4°C for 30 minutes, and the resultant supernatant (the Triton soluble fraction) was stored at −80°C. The resultant pellet was homogenized in three volumes of 50 mM Tris-HCl buffer, pH 8.0, containing 5 M guanidine-HCl plus complete protease and phosphatase inhibitors (Roche). This resuspension (the guanidine HCl soluble fraction) was incubated at 4°C for a minimum of 14 hours with shaking and was stored at −80°C. Protein concentration was determined by Bradford assay (Bio-Rad).

Samples were then analysed by human amyloid-β 6E10 Triplex (Meso Scale Discovery) following the manufacturer’s protocols. Briefly the TBS, Triton, and guanidine HCl soluble fractions were diluted into Diluent 35 (Meso Scale Discovery) and added to a precoated plate prior to addition of amyloid-β detection antibody and incubation overnight at 4°C (Meso Scale Discovery). After washing, Read Buffer (Meso Scale Discovery) was applied immediately prior to plate reading on a Meso Scale Discovery Sector Imager S600. Analyte data (pg/ml) was normalised to weight of brain region divided by total buffer homogenised in (mg/ml) to result in amount of amyloid-β analyte per brain weight (pg/mg).

### Statistical analysis

All data acquisition and analysis were undertaken blind to genotype using unique identifiers assigned at P14 and genotype codes. Mice/samples were only excluded from the study for the following technical reasons; escape from Y-maze during testing period (n = 4), failure of OCTExplorer software to correctly segment retinal layers (n = 62 layer measurements), the failure of 82E1 staining IHC because of disintegration of tissue (n =3). For Y-maze, novel object recognition, ABR, OCT and ECHO-MRI analysis, data were analysed by multi-factor ANOVA or a linear mixed model (for Y-maze in which data for specific time-points was missing because of technical issues) using sex and genotype as factors. For Home-Cage-Analysis recordings, data was analysed by linear effects mixed modelling as per Bains *et al.* (2023)^92^. Briefly, we constructed a linear mixed-effects model of distance moved as a function of the effect of sex, age and genotype. Cage ID was modelled as the random effect intercept with day of recording (timepoint) as the random effect slope. More readable model formula:

Mean activity ∼ Genotype*Age*Sex + (timepoint | Cage ID)

This model was compared to other model iterations with different combinations of sex, age and genotype with or without the interaction term and random effects structure. An ANOVA test was run to determine the statistical significance of the interaction between Age:Genotype:Sex and to inform model selection. Random effects and fixed effects found not to have a statistically significant contribution to model fit were eliminated. Models were fit using R’s “lmer” function.

For analysis of Aβ peptides and Aβ plaque coverage, additional factors of batch number were used. A Benjamini-Hochberg correction was used to control for multiple comparisons within each experiment on the cohort.

### Nanopore RNA-sequencing and data pre-processing

Total RNA was extracted from tissue samples using the Monach Total RNA kit (NEB) and assessed for quality using RNA Integrity Number (RIN) values via Tapestation (Agilent). Libraries were prepared using the PCR-cDNA Barcoding kit (SQK-PCB109) from Oxford Nanopore Technologies, following the manufacturer’s protocol. Briefly, 50 ng of total RNA were converted into cDNA through reverse transcription using Maxima H Minus Reserve Transcriptase. A strand-switching primer was added to guarantee the selection of full transcripts. cDNAs were amplified and barcoded with a unique barcode through a 15 cycle PCR with 7 minutes of extension time. Six pools composed of two samples each were created and purified using Ampure XP beads. After adapter ligation, each pool was sequenced in a PromethION flow cell (FLO-PRO002) for 72 hours. Flow cells were refuelled when the translocation speed was low and washed once and reloaded with another library aliquot. Basecalling was performed in real time with the “High accuracy basecalling” mode using Guppy 4.3.4.

### Pipeline

Quality control was performed with fastQC^93^ on the raw sequencing data. Around 30 million reads for each sample passed the initial QC step and the estimated N50 was found to be ∼1kb. The pipeline from Oxford Nanopore Technologies, IsoformSwitchAnalyzer, and PSI-Sigma tools were used to obtain transcript assembly, differential isoform usage and splicing detection, respectively (below).

### Genomic alignment

The ONT reads were aligned to the latest release of the Gencode reference genome (GRCm39) with minimap2^43^. The alignment parameters ‘-ax splice: hq -uf –secondary=no’ were used for the splice-aware alignment. The latest GENCODE annotation (GRCm39) was used for the junction information. The bam files were sorted with samtools^94^.

### Gene and isoform abundance estimation

Assembly (gtf) files of each sample were used as an input list to the *prepDE.py* function according to the manual of StringTie^95,96^. For the quantification step, all gtf files were collapsed using Stringtie --merge option. The merged gtf file was used for the quantification of the isoforms. StringTie -eB parameters were used to obtain read-count data.

### Differential expression analysis with DESeq2

Gene and isoform level differential expression analysis were performed with DESeq2^44,45^ with raw counts obtained from StringTie^96^ as described in Pertea *et al.* (2016)^84^. Sex was included to the contrast design.

### Exon and isoform usage via DEXSeq

We performed data-quality aware DTU analysis between genotypes with DEXSeq^45^. The full design formula included ∼sample + exon + Sex:exon + Genotype:exon, and tested against reduced model (∼sample + exon + Sex:exon) to detect differences due to genotype only. Then, a statistical test for differential exon usage was performed using the testForDEU function. This test compares the specified experimental design to a reduced model, providing significant usage differences across groups.

### Functional annotation via two-stage tappAS analysis

We conducted additional detailed analysis of isoform diversity and functional consequences using tappAS^48^, a Java application that integrates various functional analyses of isoforms. We used several existing tools implemented in tappAS^48^ for our analysis. To generate a functionally annotated gtf file, we used IsoAnnot_Lite_ (https://isoannot.tappas.org/) with a filtered SQANTI3^97^ gtf file obtained from the abovementioned merged gtf file from StringTie to provide input to tappAS application to perform the DTU analysis^32^. The *sqanti3_qc.py* and *sqanti3_filter.py* functions were used with the merged gtf file from gffcompare^98^. tappAS additionally outputs isoform usage and isoform switch, which was included in the GO annotations of the usage and switch analyses.

### Isoform switch analysis and predicted functional consequences

We conducted isoform switch analysis using IsoformSwitchAnalyzer^47^. Briefly, we first identified isoform switches using the merged transcript assembly from StringTie^95^, and generated isoform level count data using *prepDE.py* function. We initially checked whether our transcripts of interest had Open Reading Frames (ORF/CDS)^99,100^, and had coding potential (CPC2)^101^. Then, we have looked at whether an isoform switch results in any changes in protein domain (Pfam)^102^, whether there is any presence of signal peptides and the location of their cleavage sites using SignalP^103^, and intrinsically disordered regions (IUPred2A)^104^. Finally, we performed GO^105^ and pathway enrichment analysis on the switch events.

### Alternative splicing analysis

We used PSI-Sigma for comprehensive splicing detection for long-read RNA-seq^46^. Compared to other alternative splicing analysis methods, PSI-Sigma has several advantages over short-read splicing detection tools such as covering alternative splicing events, such as multiple-exon-skipping (MES) or more complex splicing events.

### Data Availability

All raw data will be made available via GEO, and code via GitHub.

## Software and Algorithms

|  |  |  |
| --- | --- | --- |
| Minimap2<sup>43</sup> | Long-read aligner | https://github.com/lh3/minimap2 |
| Pychopper | Identifying, orienting and trimming full-length Nanopore cDNA reads | https://github.com/epi2me-labs/pychopper |
| StringTie<sup>95</sup> | Transcriptome assembly | https://github.com/gpertea/stringtie |
| GffCompare<sup>98</sup> | Comparing and annotating transcriptome annotations | https://ccb.jhu.edu/software/stringtie/gffcompare.shtml |
| clusterProfiler<sup>95,96</sup> | Gene set enrichment analysis | https://bioconductor.org/packages/release/bioc/html/clusterProfiler.html |
| Snakemake<sup>107</sup> | Workflow management system that helps build reproducible and scalable data analyses. | https://snakemake.readthedocs.io/ |
| salmon<sup>108</sup> | Used for quantifying gene expression from RNA-seq data | https://salmon.readthedocs.io/. |
| DESeq2<sup>44</sup> | Statistical software package for differential expression analysis of RNA-seq data | https://bioconductor.org/packages/release/bioc/html/DESeq2.html |
| DEXSeq<sup>45,97</sup> | Used for analyzing differential exon usage in RNA-seq data | https://bioconductor.org/packages/release/bioc/html/DEXSeq.html |
| CoExpNets<sup>56</sup> | Weighted Gene Co-expression Network Analysis with additional k-means clustering step | https://github.com/juanbot/CoExpNets |
| SQANTI3<sup>97</sup> | Transcriptome quality control | https://github.com/ConesaLab/SQANTI |
| stageR<sup>51</sup> | Statistical analysis of gene expression data | https://bioconductor.org/packages/release/bioc/html/StageR.html |
| tappAS<sup>48,51</sup> | Computational framework for functional transcriptomics analysis | https://app.tappas.org |
| PSI-Sigma<sup>46</sup> | Splicing detection method | https://github.com/wososa/PSI-Sigma |

## Acknowledgements

This work, D.S., J.H., U.Y, N.M, F.K.W., G.B., T.L. and P.M. are supported by the UK Dementia Research Institute [award numbers UK DRI-1009 and UK DRI-1014] through UK DRI Ltd, principally funded by the Medical Research Council. D.S. also received funding from the ARUK pump priming scheme via the UCL network. J.H. and DAS are supported by the Dolby Foundation, and by the National Institute for Health Research University College London Hospitals Biomedical Research Centre. J.H. also received funding from Cure Alzheimer’s Fund. E.K.G. was also supported by the Postdoctoral Fellowship Program in Alzheimer’s Disease Research from the BrightFocus Foundation (A2021009F). R.D.U. and A.L.T. were facilitated by the Manchester NIHR Biomedical Research Centre and the Greater Manchester Local Clinical Research Network. U.Y. was funded by Ministry of Education in Turkiye. R.S.B., H.F., M.S. and S.W. were supported by Medical Research Council, Strategic Award A410-53658. F.K.W. is further supported by an Alzheimer’s Research UK Senior Research Fellowship (ARUK-SRF2018A-001). L.S. was funded by Alzheimer’s society UK and RoseTrees/Stonygate. We would like to thank Husbandry and Genotyping teams at the Mary Lyon Centre at MRC Harwell. Figure 1 drawn with BioRender.

