## Supplementary material for "Long-read transcriptomic identification of synaptic adaptation to amyloid pathology in the *App^NL-G-F^* knock-in mouse model of the earliest phase of Alzheimer’s disease": Figure S

**Running title:**

**Synaptic adaptation by altered gene splicing in amyloid mice**

**Figure S1. Amyloid-β coverage via immunohistochemistry in *App*$^{NL\text{-}G\text{-}F}$ mice.**

(A) This figure illustrates the amyloid-β coverage  in both female and male *App*$^{NL\text{-}G\text{-}F}$ mice in dorsal hippocampus and cortex compared to controls.

(B) Immunostained brain sections revealed amyloid-β staining exclusively in *App*$^{NL\text{-}G\text{-}F}$ mice, confirming the distinctive amyloid-β distribution.

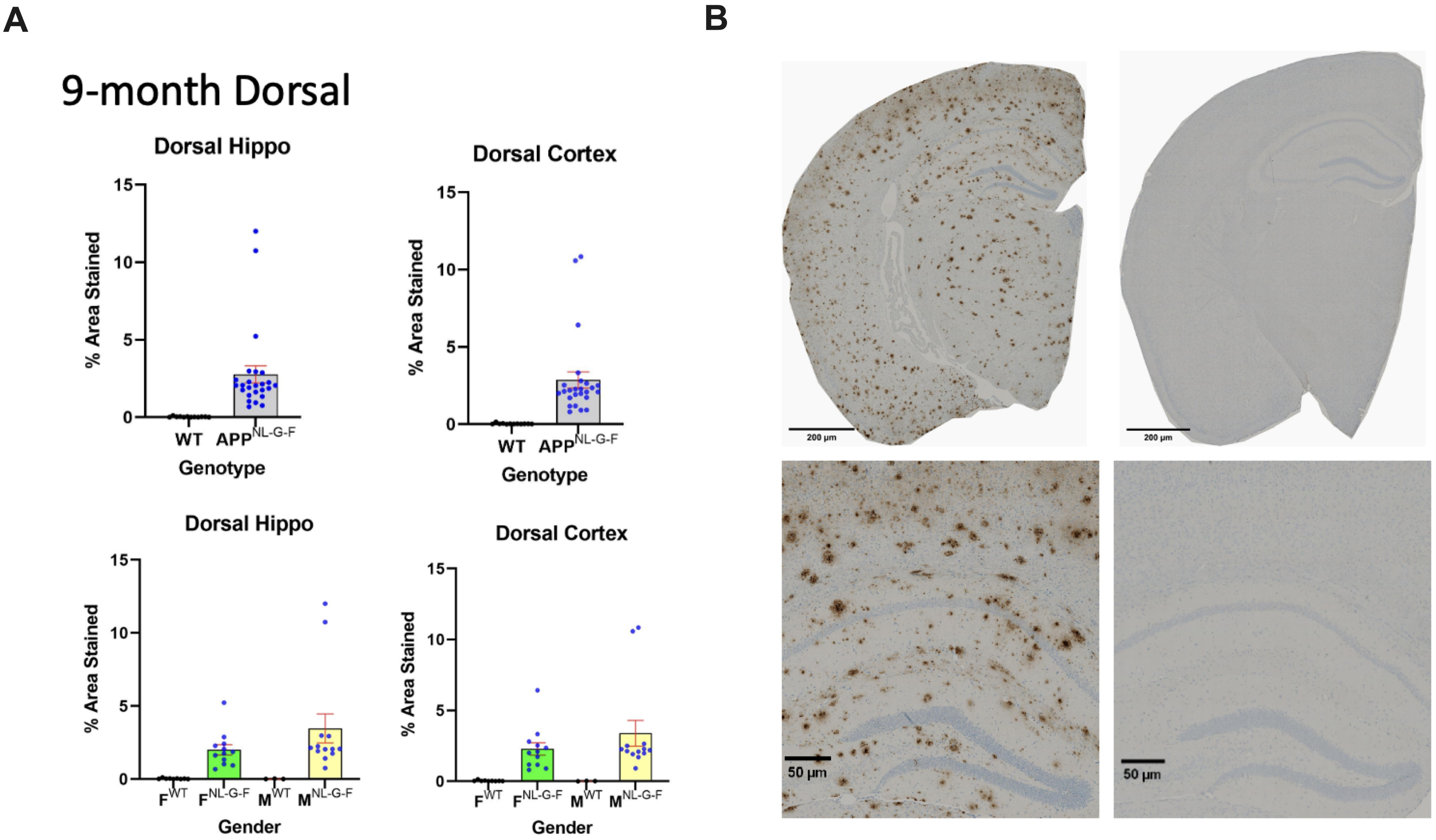


**Supplementary Figure 1**

**Figure S2. Amyloid-β coverage via immunohistochemistry in *App*$^{NL\text{-}G\text{-}F}$ mice.**

(A) This figure presents amyloid-β coverage measured by immunohistochemistry in ventral hippocampus and cortex, categorized by female and male *App*$^{NL\text{-}G\text{-}F}$ mice compared to controls. (B) Immunostained brain sections show amyloid-β staining specifically in *App*$^{NL\text{-}G\text{-}F}$ mice, confirming the amyloid-β distribution pattern in these mice.

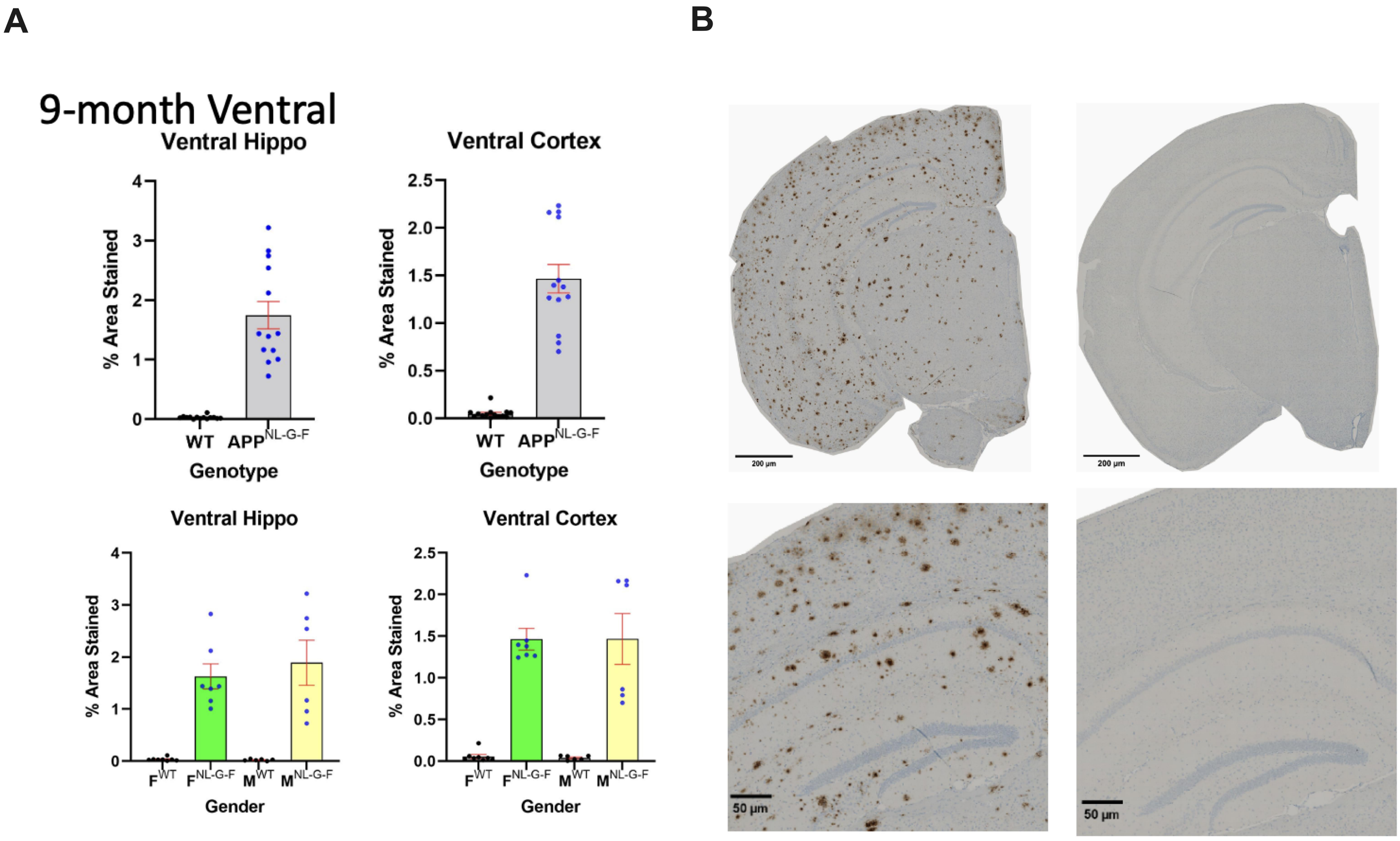


**Supplementary Figure 2**

**Figure S3. Microglial isoform-level coexpression network.**

In-depth view of the isoform-level microglial coexpression network. This figure captures isoform-isoform interactions of microglial genes to the amyloid stimulus.

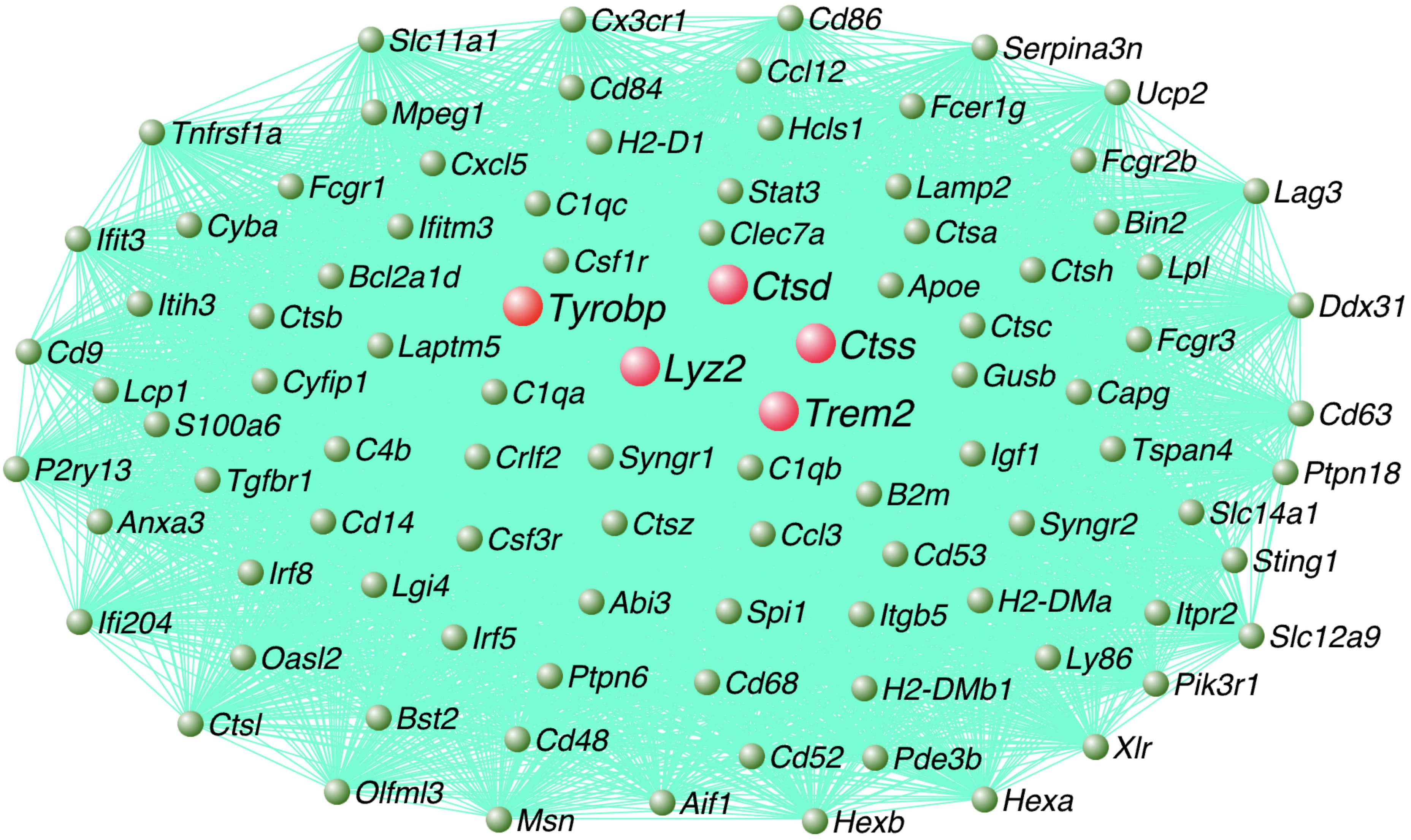


**Isoform-level microglial coexpression network in response to amyloid**

**Supplementary Figure 3**

**Figure S4. Gene ontology annotations genes with isoform switch and alternative splicing events.**

(A) Gene ontology annotations related to genes with alternative splicing events, enriched in endomembrane system organizations and actin-based pathways, including actin filament binding and actin-based cell projections.

(B) Gene ontology annotations highlight that genes exhibiting isoform switches are predominantly enriched in developmental pathways, protein binding, and cell junctions.

A

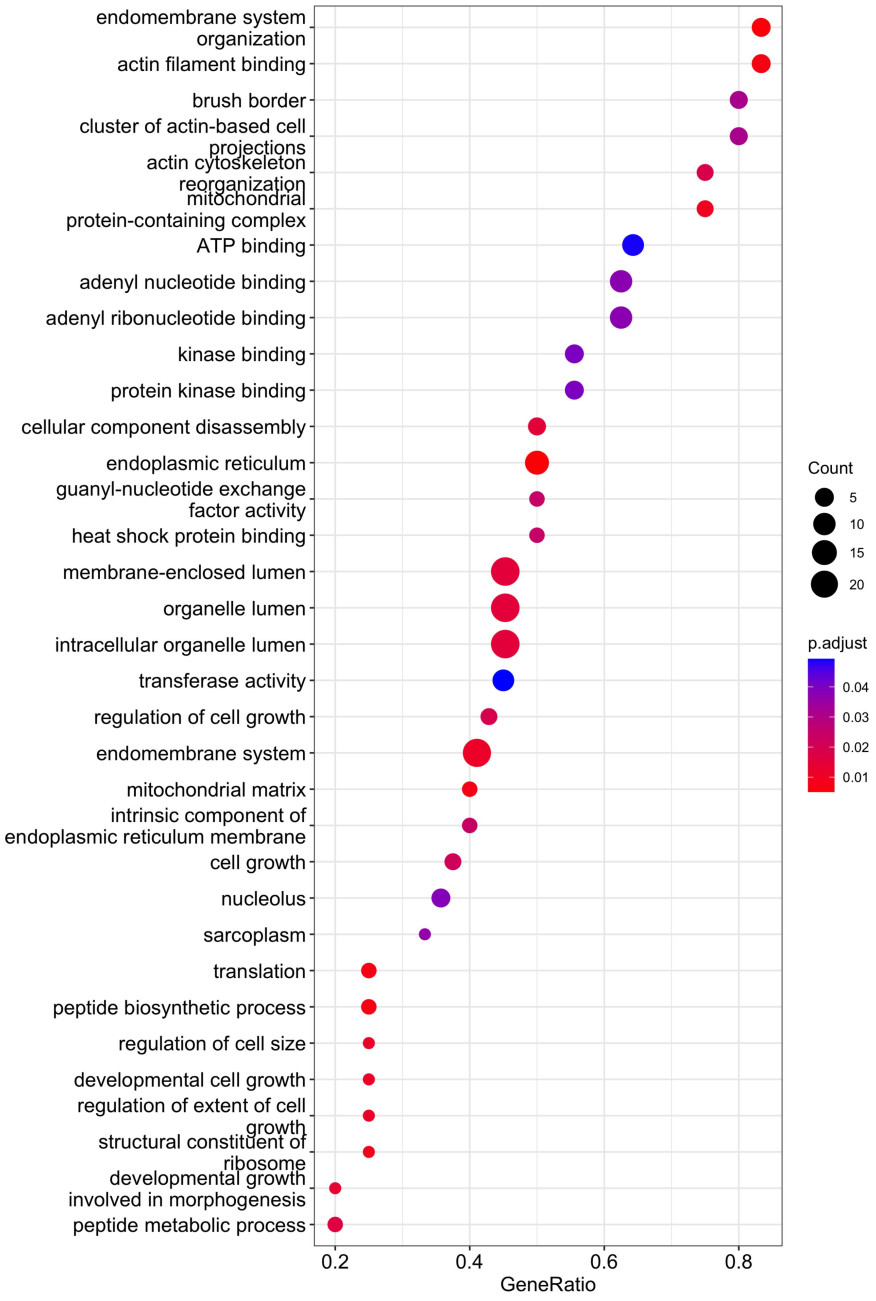


**Gene ontology annotation of spliced genes**

B

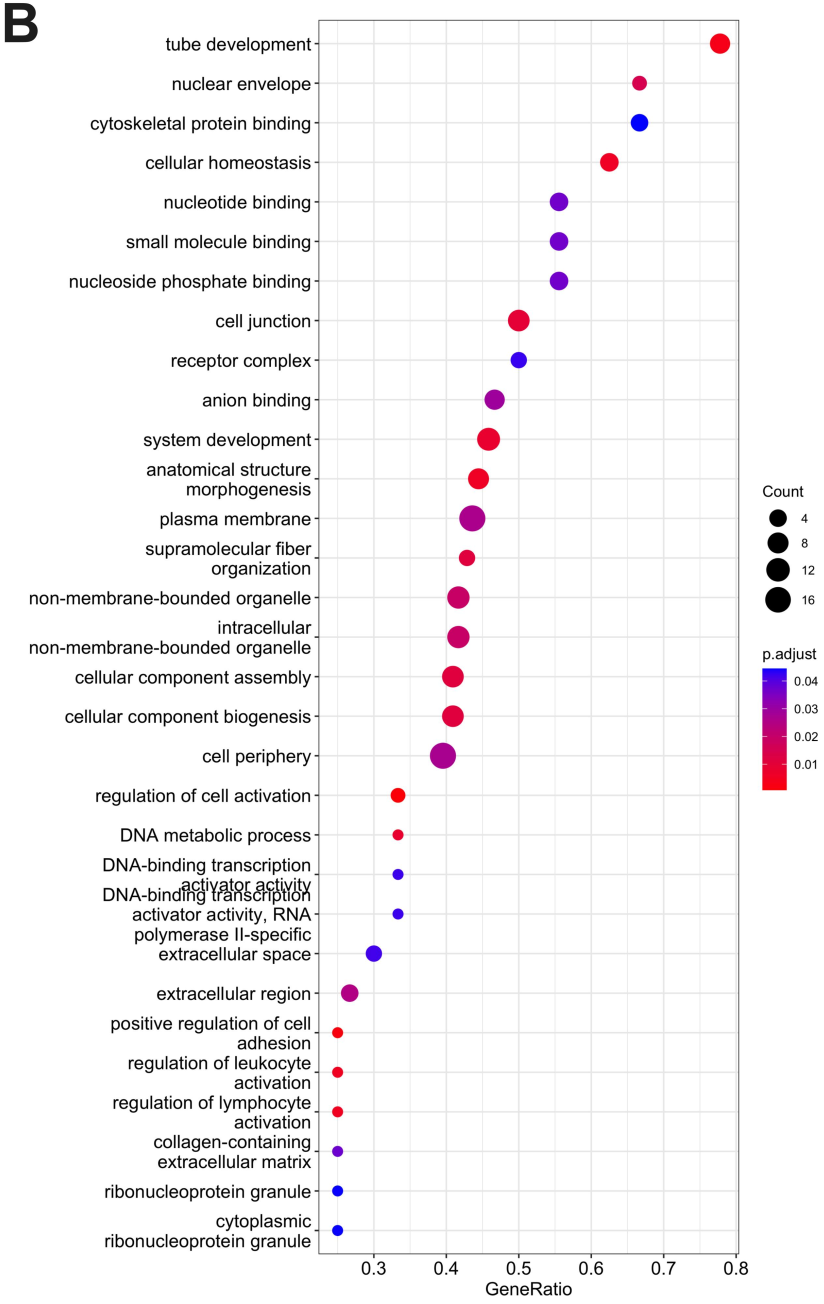


**Gene ontology annotation of switched genes**

### Supplementary Figure 4
