## Supplementary material for "Long-read transcriptomic identification of synaptic adaptation to amyloid pathology in the *App^NL-G-F^* knock-in mouse model of the earliest phase of Alzheimer’s disease": Table S2

Table S2. Home-cage data during dark, light and transition phase of daily cycles.

| Time period of analysis | Genotype | Age | Sex | Mean (mm) | Standard Error | Degrees of freedom | Age comparison | Sex comparison | Lighting phase comparison | Genotype comparison | p |
| --- | --- | --- | --- | --- | --- | --- | --- | --- | --- | --- | --- |
| Complete dark phase | Hom | 10_weeks | f | 23.97507192 | 1.118050477 | 14.52337677 | 10_weeks | f | Dark | Hom - WT | 0.709985 |
| Complete dark phase | WT | 10_weeks | f | 25.20288021 | 1.171209739 | 16.48748326 | 32_weeks | f | Dark | Hom - WT | 0.709985 |
| Complete dark phase | Hom | 32_weeks | f | 24.6447706 | 1.137629139 | 15.53328908 | 10_weeks | m | Dark | Hom - WT | 0.349296 |
| Complete dark phase | WT | 32_weeks | f | 23.39297606 | 1.171209739 | 16.48748326 | 32_weeks | m | Dark | Hom - WT | 0.709985 |
| Complete dark phase | Hom | 10_weeks | m | 22.60142444 | 1.171209739 | 16.48748326 | 10_weeks | f | Light | Hom - WT | 0.762964 |
| Complete dark phase | WT | 10_weeks | m | 19.91038134 | 1.327086087 | 22.01433182 | 32_weeks | f | Light | Hom - WT | 0.836803 |
| Complete dark phase | Hom | 32_weeks | m | 18.35822843 | 1.230062787 | 19.9071744 | 10_weeks | m | Light | Hom - WT | 0.349296 |
| Complete dark phase | WT | 32_weeks | m | 19.60554201 | 1.372885434 | 25.03998661 | 32_weeks | m | Light | Hom - WT | 0.836803 |
| Complete light phase | Hom | 10_weeks | f | 17.68307053 | 1.118050477 | 14.52337677 | 10_weeks | f | Dark | Hom - WT | 0.709985 |
| Complete light phase | WT | 10_weeks | f | 18.45360016 | 1.171209739 | 16.48748326 | 32_weeks | f | Dark | Hom - WT | 0.709985 |
| Complete light phase | Hom | 32_weeks | f | 17.16004618 | 1.137629139 | 15.53328908 | 10_weeks | m | Dark | Hom - WT | 0.349296 |
| Complete light phase | WT | 32_weeks | f | 16.87632119 | 1.171209739 | 16.48748326 | 32_weeks | m | Dark | Hom - WT | 0.709985 |
| Complete light phase | Hom | 10_weeks | m | 18.63021713 | 1.171209739 | 16.48748326 | 10_weeks | f | Light | Hom - WT | 0.762964 |
| Complete light phase | WT | 10_weeks | m | 15.58131841 | 1.327086087 | 22.01433182 | 32_weeks | f | Light | Hom - WT | 0.836803 |
| Complete light phase | Hom | 32_weeks | m | 15.48434529 | 1.230062787 | 19.9071744 | 10_weeks | m | Light | Hom - WT | 0.349296 |
| Last 30 minutes of dark phase | Hom | 10_weeks | f | 166.3175772 | 28.66411545 | 86.46601919 | 10_weeks | f | na | Hom - WT | 0.864951 |
| Last 30 minutes of dark phase | WT | 10_weeks | f | 134.7965778 | 28.66411545 | 86.46601919 | 32_weeks | f | na | Hom - WT | 0.864951 |
| Last 30 minutes of dark phase | Hom | 32_weeks | f | 160.5428487 | 14.35564854 | 24.08660075 | 10_weeks | m | na | Hom - WT | 0.906574 |
| Last 30 minutes of dark phase | WT | 32_weeks | f | 143.4817911 | 14.35564854 | 24.08660075 | 32_weeks | m | na | Hom - WT | 0.864951 |
| Last 30 minutes of dark phase | Hom | 10_weeks | m | 93.39070978 | 15.89941733 | 36.37693631 | 10_weeks | f | na | Hom - WT | 0.864951 |
| Last 30 minutes of dark phase | WT | 10_weeks | m | 90.58583731 | 17.61010764 | 33.405285 | 32_weeks | f | na | Hom - WT | 0.864951 |
| Last 30 minutes of dark phase | Hom | 32_weeks | m | 87.2516221 | 15.77255835 | 34.51616023 | 10_weeks | m | na | Hom - WT | 0.906574 |
| Last 30 minutes of dark phase | WT | 32_weeks | m | 98.11778327 | 17.61010764 | 33.405285 | 32_weeks | m | na | Hom - WT | 0.864951 |
| First 30 minutes of dark phase | Hom | 10_weeks | f | 323.0711575 | 27.26600156 | 32.83807996 | 10_weeks | f | na | Hom - WT | 0.672825 |
| First 30 minutes of dark phase | WT | 10_weeks | f | 360.4154358 | 29.14858175 | 32.83807996 | 32_weeks | f | na | Hom - WT | 0.71991 |
| First 30 minutes of dark phase | Hom | 32_weeks | f | 273.8131486 | 28.48819262 | 37.05562601 | 10_weeks | m | na | Hom - WT | 0.110161 |
| First 30 minutes of dark phase | WT | 32_weeks | f | 259.079093 | 29.14858175 | 32.83807996 | 32_weeks | m | na | Hom - WT | 0.672825 |
| First 30 minutes of dark phase | Hom | 10_weeks | m | 324.9819778 | 29.14858175 | 32.83807996 | 10_weeks | f | na | Hom - WT | 0.672825 |
| First 30 minutes of dark phase | WT | 10_weeks | m | 220.8320681 | 34.48906704 | 32.83807996 | 32_weeks | f | na | Hom - WT | 0.71991 |
| First 30 minutes of dark phase | Hom | 32_weeks | m | 218.3467793 | 32.67560416 | 44.49039706 | 10_weeks | m | na | Hom - WT | 0.110161 |
| First 30 minutes of dark phase | WT | 32_weeks | m | 185.0474001 | 37.1523098 | 40.18549601 | 32_weeks | m | na | Hom - WT | 0.672825 |
