## Supplementary material for "Long-read transcriptomic identification of synaptic adaptation to amyloid pathology in the *App^NL-G-F^* knock-in mouse model of the earliest phase of Alzheimer’s disease": Table S3

Table S3: Sensory Phenotyping.

|  |  |  | Optical Coherence Tomography | | | | | | | | | | Auditory Brainstem Response | | | | | |
| --- | --- | --- | --- | --- | --- | --- | --- | --- | --- | --- | --- | --- | --- | --- | --- | --- | --- | --- |
|  |  |  | RNFL | GCL | IPL | INL | OPL | ONL | IS OS | Outer segment | OPR Subretinal virtual space | RPE | Click | 6Hz | 12Hz | 18Hz | 24Hz | 30Hz |
| Female | Young | Wildtype | 17.76 ±0.47 | 16.77 ±0.55 | 28.93 ±0.55 | 20.26 ±0.29 | 18.34 ±0.37 | 79.78 ±0.78 | 11.27 ±0.22 | 25.63 ±1.27 | 28.40 ±1.19 | 12.44 ±0.03 | 43.33 ±2.07 | 47.92 ±3.23 | 46.25 ±3.15 | 47.5 ±3.05 | 50.42 ±3.87 | 53.75 ±5.91 |
|  |  | APP^NL-G-F/NL-G-F^ | 18.97 ±0.9 | 15.92 ±0.52 | 28.53 ±0.42 | 18.85 ±0.46 | 18.33 ±0.29 | 78.94 ±1.02 | 11.21 ±0.2 | 22.9 ±1.58 | 30.4 ±1.59 | 12.46 ±0.02 | 37.92 ±2.57 | 48.33 ±3.45 | 44.58 ±3.17 | 40.83 ±2.81 | 45 ±4.31 | 50.42 ±3.11 |
|  | Late-middle age | Wildtype | 19.34 ±0.75 | 18.13 ±0.29 | 28.43 ±0.34 | 18.93 ±0.37 | 18.52 ±0.21 | 79.37 ±0.53 | 11.45 ±0.15 | 26.83 ±0.6 | 36.39 ±0.93 | 12.36 ±0.02 |  |  |  |  |  |  |
|  |  | APP^NL-G-F/NL-G-F^ | 19.95 ±0.62 | 15.78 ±0.46 | 28.71 ±0.51 | 18.5 ±0.39 | 18.57 ±0.2 | 78.85 ±0.75 | 11.2 ±0.21 | 23.78 ±1.04 | 38.01 ±1.17 | 12.37 ±0.01 |  |  |  |  |  |  |
| Male | Young | Wildtype | 19.58 ±0.74 | 16.72 ±0.41 | 28.47 ±0.47 | 19.21 ±0.48 | 18.63 ±0.21 | 78.25 ±0.68 | 11.11 ±0.12 | 24.05 ±1.19 | 33.56 ±1.39 | 12.43 ±0.02 | 33.33 ±4.04 | 42.5 ±3.82 | 34.17 ±3.52 | 33.33 ±3.8 | 43.33 ±3.8 | 50.83 ±5.07 |
|  |  | APP^NL-G-F/NL-G-F^ | 19.74 ±0.86 | 15.2 ±0.5 | 28.78 ±0.37 | 18.25 ±0.38 | 18.54 ±0.27 | 78.66 ±1.17 | 11.04 ±0.12 | 23.83 ±1.8 | 33.33 ±1.56 | 12.46 ±0.02 | 35.83 ±2.67 | 48.75 ±2.39 | 37.5 ±3.15 | 38.33 ±2.25 | 45.83 ±1.83 | 50 ±5.91 |
|  | Late-middle age | Wildtype | 20.1 ±0.67 | 15.62 ±0.37 | 28.22 ±0.49 | 18.59 ±0.43 | 18.1 ±0.25 | 80.46 ±0.78 | 11.13 ±0.16 | 26.06 ±1.07 | 37.75 ±1.03 | 12.36 ±0.02 |  |  |  |  |  |  |
|  |  | APP^NL-G-F/NL-G-F^ | 19.36 ±0.74 | 16.2 ±0.54 | 28.43 ±0.47 | 18.38 ±0.49 | 17.91 ±0.33 | 78.46 ±1.06 | 10.77 ±0.13 | 27.33 ±0.9 | 35.94 ±0.89 | 12.37 ±0.02 |  |  |  |  |  |  |
| Repeat Measures ANOVA analysis | | Genotype | 0.466 | 0.015 | 0.609 | 0.036 | 0.972 | 0.235 | 0.109 | 0.1795 | 0.660 | 0.177 | 0.611 | 0.328 | 0.790 | 0.785 | 0.704 | 0.666 |
|  |  | Genotype X Age | 0.721 | 0.401 | 0.493 | 0.110 | 0.698 | 0.274 | 0.653 | 0.598 | 0.918 | 0.501 |  |  |  |  |  |  |
|  |  | Genotype X Sex | 0.294 | 0.035 | 0.466 | 0.440 | 0.696 | 0.774 | 0.913 | 0.149 | 0.372 | 0.767 | 0.173 | 0.391 | 0.426 | 0.062 | 0.305 | 0.795 |
|  |  | Genotype X Age X Sex | 0.745 | 0.028 | 0.489 | 0.788 | 0.847 | 0.244 | 0.889 | 0.555 | 0.005 | 0.725 |  |  |  |  |  |  |
|  |  | Age | 0.444 | 0.453 | 0.526 | 0.123 | 0.457 | 0.722 | 0.676 | 0.0315 | 0.001 | 0.004 |  |  |  |  |  |  |
