## Supplementary material for "Long-read transcriptomic identification of synaptic adaptation to amyloid pathology in the *App^NL-G-F^* knock-in mouse model of the earliest phase of Alzheimer’s disease": Table S4

Table S4: ECHO-MRI data and statistical analysis.

|  |  |  | Body weight | Lean mass |
| --- | --- | --- | --- | --- |
| Female data | Young | Wildtype | 20.59 ±0.26 | 13.03 ±0.26 |
|  |  | APP^NL-G-F/NL-G-F^ | 20.45 ±0.26 | 12.93 ±0.23 |
|  | Middle age | Wildtype | 23.88 ±0.26 | 15.22 ±0.25 |
|  |  | APP^NL-G-F/NL-G-F^ | 23.61 ±0.36 | 15.33 ±0.26 |
|  | Late-middle age | Wildtype | 28.79 ±0.73 | 15.53 ±0.73 |
|  |  | APP^NL-G-F/NL-G-F^ | 27.06 ±0.58 | 15.49 ±0.29 |
| Male data | Young | Wildtype | 26.16 ±0.29 | 18.02 ±0.29 |
|  |  | APP^NL-G-F/NL-G-F^ | 26.07 ±0.29 | 18.29 ±0.28 |
|  | Middle age | Wildtype | 30.52 ±0.31 | 21.29 ±0.32 |
|  |  | APP^NL-G-F/NL-G-F^ | 30.75 ±0.39 | 22.39 ±0.33 |
|  | Late-middle age | Wildtype | 34.7 ±0.61 | 21.48 ±0.59 |
|  |  | APP^NL-G-F/NL-G-F^ | 34.74 ±0.75 | 22.16 ±0.35 |
| Repeat Measures ANOVA analysis (p values) | Genotype | | 0.2731 | 0.2593 |
|  | Genotype X Age | | 0.1197 | 0.2321 |
|  | Genotype X Sex | | 0.4043 | 0.2459 |
|  | Genotype X Age X Sex | | 0.4894 | 0.5652 |
|  | Age | | < 0.0001 | < 0.0001 |
|  | Sex | | < 0.0001 | < 0.0001 |
|  | Age X Sex | | 0.0428 | 0.0001 |
