## Supplementary material for "Long-read transcriptomic identification of synaptic adaptation to amyloid pathology in the *App^NL-G-F^* knock-in mouse model of the earliest phase of Alzheimer’s disease": Table S1

Table S1. Behavioural phenotyping with forced Y-maze and novel object recognition.

|  |  |  | Forced Y-maze | | | Novel object recognition | | |
| --- | --- | --- | --- | --- | --- | --- | --- | --- |
|  |  |  | Duration Novelty Preference Ratio | Number of Entries Novelty Preference Ratio | Distance moved | Duration Novelty Preference Ratio | Number of Investigations Novelty Preference Ratio | Distance moved |
| Female | Young | Wildtype | 0.581 ±0.023 | 0.595 ±0.047 | 5134 ±272 | 0.674 ±0.036 | 0.613 ±0.027 | 1612 ±145 |
|  |  | APP^NL-G-F/NL-G-F^ | 0.569 ±0.0.39 | 0.583 ±0.024 | 5246 ±411 | 0.621 ±0.032 | 0.579 ±0.026 | 1800 ±187 |
|  | Late-middle age | Wildtype | 0.63 ±0.05 | 0.546 ±0.028 | 4063 ±229 | 0.659 ±0.027 | 0.621 ±0.017 | 1653 ±107 |
|  |  | APP^NL-G-F/NL-G-F^ | 0.422 ±0.064 | 0.451 ±0.041 | 3884 ±322 | 0.676 ±0.023 | 0.606 ±0.017 | 1760 ±72 |
| Male | Young | Wildtype | 0.568 ±0.089 | 0.531 ±0.07 | 5396 ±307 | 0.659 ±0.029 | 0.608 ±0.013 | 1955 ±103 |
|  |  | APP^NL-G-F/NL-G-F^ | 0.664 ±0.047 | 0.593 ±0.037 | 4988 ±301 | 0.699 ±0.038 | 0.607 ±0.027 | 2195 ±157 |
|  | Late-middle age | Wildtype | 0.686 ±0.034 | 0.596 ±0.027 | 4766 ±254 | 0.638 ±0.027 | 0.613 ±0.024 | 1858 ±113 |
|  |  | APP^NL-G-F/NL-G-F^ | 0.591 ±0.048 | 0.524 ±0.032 | 3844 ±227 | 0.739 ±0.029 | 0.652 ±0.027 | 1679 ±90 |
| Repeat Measures ANOVA analysis | | Genotype | 0.161 | 0.223 | 0.154 | 0.135 | 0.916 | 0.364 |
|  |  | Genotype X Age | 0.012 | 0.048 | 0.126 | 0.069 | 0.202 | 0.146 |
|  |  | Genotype X Sex | 0.154 | 0.298 | 0.19 | 0.070 | 0.287 | 0.551 |
|  |  | Genotype X Age X Sex | 0.958 | 0.509 | 0.896 | 0.955 | 0.724 | 0.325 |
|  |  | Age | 0.641 | 0.16 | < 0.0001 | 0.283 | 0.386 | 0.078 |
|  |  | Sex | 0.079 | 0.304 | 0.588 | 0.418 | 0.141 | 0.032 |
|  |  | Age X Sex | 0.277 | 0.187 | 0.162 | 0.844 | 0.833 | 0.077 |
